# Mettl14-mediated m^6^A modification ensures the cell cycle progression of late-born retinal progenitor cells

**DOI:** 10.1101/2022.06.11.495708

**Authors:** Liang Li, Yue Sun, Alexander E. Davis, Man-Ru Wu, Cheng-Hui Lin, Jun B. Ding, Sui Wang

## Abstract

Neural progenitor cells lengthen their cell cycle to prime themselves for differentiation as development proceeds. It is currently not clear how they counter this lengthening and avoid being halted in the cell cycle. Here, we show that m^6^A (N^6^-methyladenosine) methylation of cell cycle-related mRNAs ensures the proper cell cycle progression of late-born retinal progenitor cells (RPCs), which are born towards the end of retinogenesis and have long cell cycle durations. Conditional deletion of *Mettl14*, which is required for depositing m^6^A on mRNAs, significantly reduced the level of m^6^A modification in the developing mouse retina. This led to delayed cell cycle exit of late-born RPCs and neuronal cell death in mature retina, but appeared to have no effect on retinal development prior to birth. M^6^A-seq, which maps m^6^A modified mRNAs, and single cell transcriptomic analyses revealed that mRNAs involved in slowing down cell cycle progression were highly enriched for m^6^A modification, which could target them for degradation and guarantee proper cell cycle progression of late-born RPCs. In addition, we identified *Zfp292* as a novel target of m^6^A and potent inhibitor of RPC cell cycle progression. Overall, our work establishes m^6^A modification as an important mechanism countering cell cycle lengthening in late-born retinal progenitor cells.

## Introduction

During central nervous system (CNS) development, the cell cycle progression and differentiation of neural progenitor cells are precisely coordinated (Hardwick and Philpott 2014; Hardwick et al. 2015; Homem et al. 2015; Miles and Tropepe 2016; Belmonte-Mateos and Pujades 2022). One of the common observations across different cell lineages is that neural progenitor cells lengthen their cell cycles as development proceeds, which was found to be necessary for accumulating differentiation related factors (Hardwick and Philpott 2014; Miles and Tropepe 2016). While studies have focused on understanding cell cycle lengthening and its association with differentiation, it is not clear how neural progenitors counter the lengthening and avoid being halted within the cell cycle for an extended time. An improved understanding of how neural progenitor cells control cell cycle progression has broad implications and can benefit cancer and regeneration-based studies.

The neural retina is a highly advantageous model for studying the cell cycle progression of neural progenitor cells. Access to retinal progenitor cells (RPCs) using techniques such as in vivo electroporation and gamma-retrovirus based lineage tracing provides a strong platform for studying RPC cell cycle progression and differentiation (Matsuda and Cepko 2007; Turner and Cepko 1987). The length of the RPC cell cycle increases during retinal development from approximately 14 hours at embryonic day 14 (E14) to about 30 hours at neonatal stages in rodent models (Alexiades and Cepko 1996). This cell cycle lengthening coincides with RPCs transitioning from early-born RPCs that differentiate into early-born neurons, such as ganglion cells and cones, to late-born RPCs that adopt late-born cell fates, including bipolar interneurons and Müller glial cells (Cepko et al. 1996). While the exact mechanisms controlling the lengthening of the RPC cell cycle have yet to be elucidated, cell cycle components, including cyclin, CDK and CDK inhibitors, transcription factors, such as Vsx2, Pax6, Meis1/2 and Six3/6, epigenetic regulators, and differentiation related genes all play important roles (Miles and Tropepe 2016). RPCs must counteract this cell cycle lengthening to ensure proper cell cycle progression. However, not much is known about the mechanisms that deter cell cycle lengthening, which is especially relevant to late-born RPCs with extended cell cycle duration.

Post-transcriptional m^6^A (N^6^-methyladenosine) modification is the most common internal modification on eukaryotic mRNAs and has been shown to play important roles in numerous biological processes, including neural development (Yu et al. 2021). It regulates many aspects of mRNA metabolism, including stability, translation, splicing and nuclear export (Jiang et al. 2021). m^6^A is deposited by the “writer” complex, which contains the core components Mettl3 (Methyltransferase like-3), Mettl14 (Methyltransferase like-14) and WTAP (Wilms Tumor 1-Associating Protein) (Yang et al. 2018). Deleting either *Mettl3* or *Mettl14* can deplete m^6^A. Yoon et al. 2017 showed that m^6^A depletion resulted in a prolonged cell cycle in neural stem cells in the cerebral cortex and extended neurogenesis in postnatal stages in *Mettl14^f/f^;Nestin-Cre* mice. Using the same animal model, Wang et al. 2018 reported that depletion of m^6^A in neural stem cells led to decreased proliferation and premature differentiation in the cortex. Even though additional studies may be needed to explain the discrepancy, these studies strongly support the importance of m^6^A in regulating the cell cycles of neural progenitor cells and encouraged us to examine the function of m^6^A using RPC as a model system.

Here we show that m^6^A methylation on mRNAs is a key mechanism that enables late-born RPCs to counter cell cycle lengthening. We conditionally deleted *Mettl14* in RPCs from the beginning of retinogenesis and found that m^6^A modification ensures proper cell cycle progression of late-born RPCs.

## Results

### Depletion of Mettl14-mediated m^6^A modification induces retinal developmental delay at neonatal stages and neuronal cell death in differentiated retina

The genes involved in regulating m^6^A dynamics are robustly expressed in the developing retina as revealed by quantitative PCR (qPCR) (Fig S1A&B). We detected the METTL14 protein, a required component of the m^6^A writer complex, in the retina starting from embryonic day 12.5 (E12.5) to adulthood in most retinal cell types (Figure S1C&E). To study the function of m^6^A in the retina, we generated a *Mettl14* conditional knockout mice (Mettl14-cKO) by breeding *Mettl14^f/f^* mice with the *Chx10-Egfp/Cre* mouse line, which expresses EGFP and Cre simultaneously starting from approximately E13.5 in RPCs (Figure S1F) (Rowan and Cepko 2004). We confirmed that the Mettl14 protein level was significantly reduced at embryonic (E15.5) and neonatal stages (P0) in Mettl14-cKO retinas (*Mettl14^f/f^;Chx10-Egfp/Cre*) compared to controls (*Mettl14^+/+^;Chx10-Egfp/Cre*) (Figure S1C). Total m^6^A methylation levels were also significantly lower in Mettl14-cKO retinas compared to controls (Figure S1D), demonstrating successful depletion of Mettl14-mediated m^6^A modification in the retina.

We carefully examined the retinas of Mettl14-cKO mice at different developmental stages and in adults. Prior to birth, we did not observe any significant defects in Mettl14-cKO retinas at E13.5 and E15.5 (Figure S2A-C). The number of Otx2 positive photoreceptor precursors and Brn3a positive developing ganglion cells was not significantly different in Mettl14-cKO and control retinas (Figure S2D-H). At P0, the retinal thickness and the number of RPCs stayed comparable between Mettl14-cKO and control retinas (Figure 1A&E and Figure S2I&J). Therefore, Mettl14-mediated m^6^A modification appears to be dispensable for early retinal development.

At neonatal stages, most RPCs finish dividing by P7 in control retinas. We can observe nicely formed outer plexiform layer (OPL) at P7 (Figure 1 B1-2). In Mettl14-cKO retinas, while the retinal thickness was unaffected (Figure 1E), the outer plexiform layer (OPL) was not well formed (Figure 1 B3-5, yellow arrow). We also observed the appearance of rosette-like structure in the central retina, suggesting defects in retinal development (Figure 1 B3&B5, while arrows).

**Figure 1.**
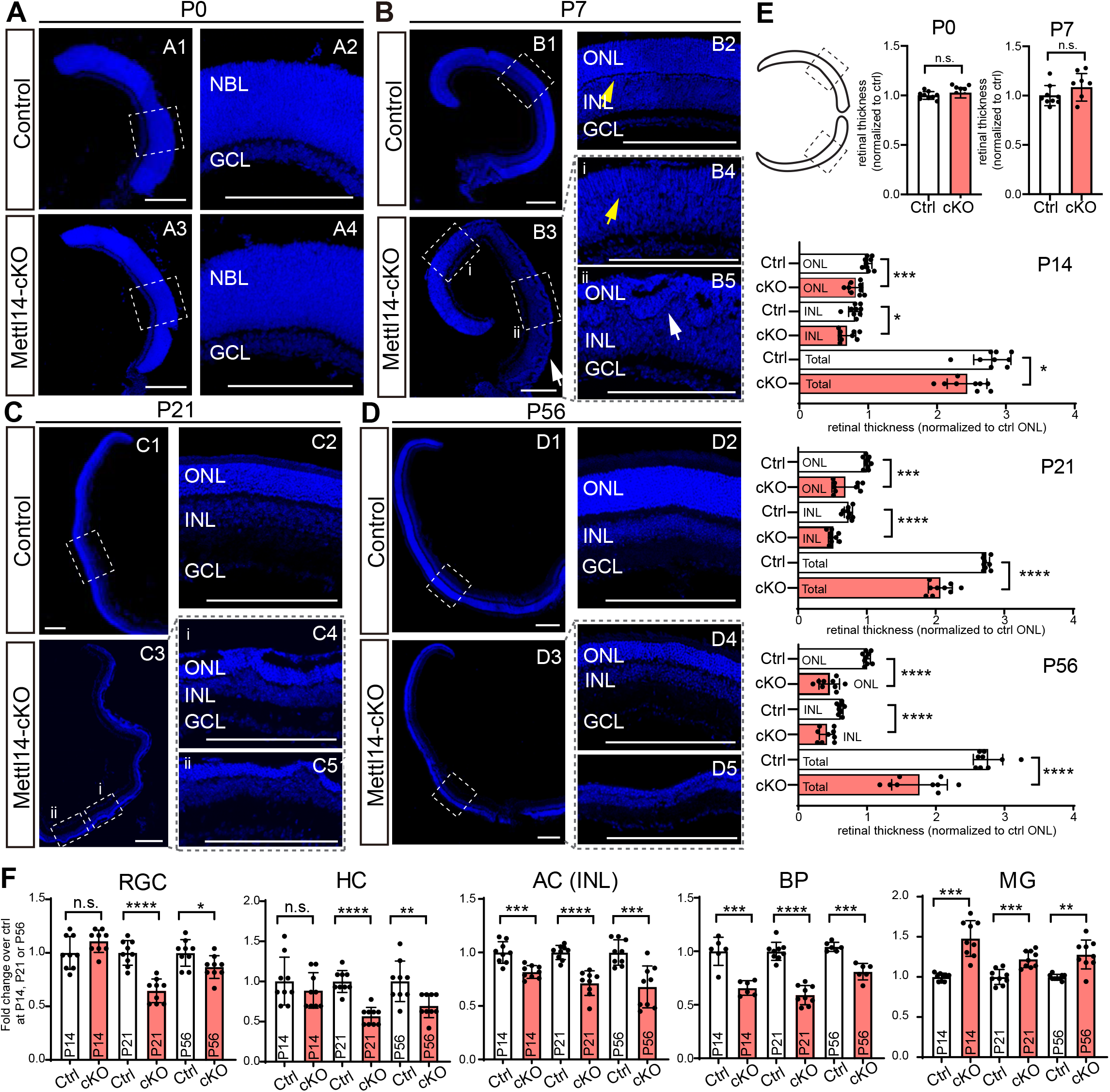
Depletion of Mettll4 leads to retinal developmental defects at neonatal stages and neuronal cell death in differentiated retina. **(A-D)** DAPI staining showing the gross morphology of the control and Mettl14-cKO retinas at P0, P7, P21 and P56. Yellow arrows in B point to the position of outer plexiform layer (OPL). White arrows in B point to the rosette-like structures. NBL: neuroblast layer; A2, A4, B2, B4-5, C2, C4-5, D2 and D4 are higher magnification views of the highlighted regions in A1, A3, B1, B3, C1, C3, D1 and D3 respectively. D5 is the higher magnification view of the highlighted region in Figure S1H. ONL: outer nuclear layer; INL: inner nuclear layer; IPL: inner plexiform layer; GCL: ganglion cell layer. Scale bar: 200μm. **(E)** The relative retinal thickness of Mettl14-cKO and control retinas at P0, P7, P14, P21 and P56 (normalized to Ctrl ONL). The highlighted regions in the cartoon are the regions we quantified (central retina). Mean ± SD (n=6-9; *P<0.05, ***P< 0.001; ****p< 0.0001; Twotailed Student’s T test). ONL: the thickness of the ONL; INL: the thickness of the INL; Total: the thickness of the entire retina. Ctrl: control retina; cKO: Mettl14-cKO retina. **(F)** The number of different retinal cell types in control and Mettl14-cKO retinas at P14, P21 and P56. Mean ± SD (n=9; *P<0.05, **P<0.01, ***P< 0.001; ****P< 0.0001; Two-tailed Student’s T test). RGC: retinal ganglion cells; HC: horizontal cells; AC (INL): amacrine cells in INL; BP: bipolar cells; MG: Müller glial cells. Ctrl: control retina; cKO: Mettl14-cKO retinas.

Retinal development completes at around P21 in mice. At P21, Mettl14-cKO mice showed significantly decreased retinal thickness compared to those of controls (Figure 1C&E). The rosette-like structure spread to the peripheral regions of the retina and disappeared from the central regions (Figure 1 C3-5). In mature retinas (P56), the size of Mettl14-cKO eyeball was significantly reduced (Figure S1G) and the retina was even thinner compared to controls (Figure 1D&E). This was companied by the disappear of the rosette-like structure. In approximately 30% of the Mettl14-cKO retinas, the outer nuclear layer (where photoreceptors reside) only contained a few rows of cells (Figure 1D5 and Figure S1H). These results suggest that m6A depletion also affects the survival of mature retinal cells.

To further elucidate the developmental defects of Mettl14-cKO retinas at around P7, we stained the retinas with Bassoon antibody, which is a well-established marker for ribbon synapses residing in the OPL (Figure 2A) (Dick et al. 2003). The absence of Bassoon expression in Mettl14-cKO further confirmed the delayed formation of OPL (Figure 2B). In addition, most RPCs exit cell cycle and undergo differentiation at P7 in control retinas (Figure 2C). However, we observed a significant number of Ki67 (cell cycle marker) positive RPCs in the central and mid-peripheral areas of Mettl14-cKO retinas at P7, while Ki67+ RPCs were rarely found in controls in these areas at this stage (Figure 1E). These P7 Ki67+ cells incorporated the thymidine analog EdU (5-ethynyl-2’-deoxyuridine), which labels cells in the S phase of the cell cycle (Fig 1F&G) (Buck et al. 2008), further supporting their identities as RPCs. Most Mettl14-cKO RPCs exited cell cycle at around P10 (Figure 2D). Moreover, we examined the cell death of retinal cells during development in Mettl14-cKO and control retinas. The number of caspase 3 positive cells peaked at P5 and P9 in controls, which was the naturally occurring cell death during development. These peaks were delayed to P6 and P10 in Mettl14-cKO retinas (Figure 2E). Taken together, these results support that the retinal development is delayed at neonatal stages in the absence of *Mettl14.*

**Figure 2.**
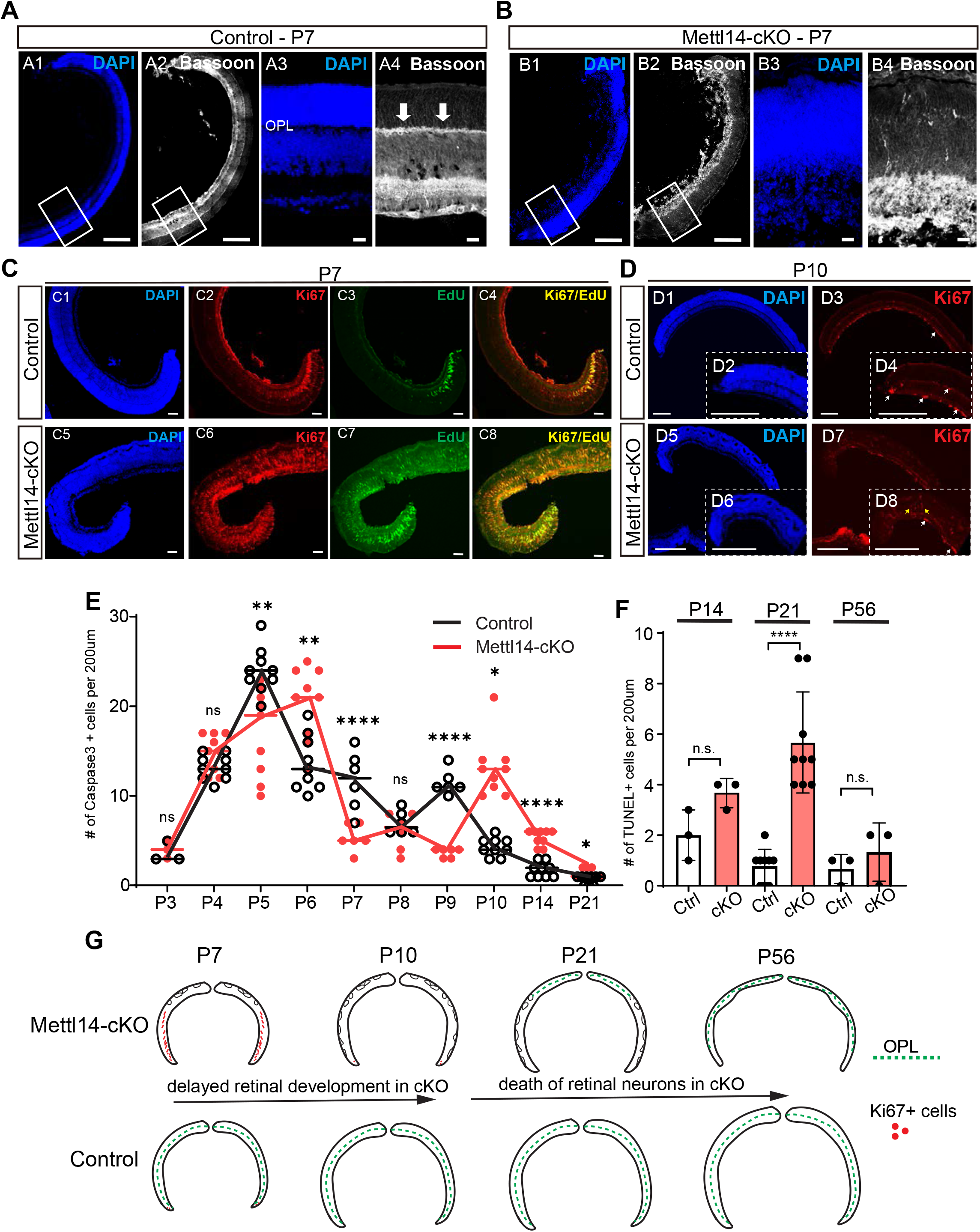
Mettl14-cKO retinas contain excessive RPCs at P7 and show significant neuronal cell death in adults. **(A-B)** Bassoon staining of control and Mettl14-cKO retinas at P7. White arrows: Bassoon staining showing OPL. A3 & A4, or B3 & B4 are higher magnification views of the highlighted regions in A1 & A2 or B1 & B2. OPL: outer plexiform later. Scale bar: 200μm (A1, A2, B1 and B2) and 20μm (A3, A4, B3 and B4). **(C)** Ki67+ and EdU+ cells in P7 control and Mettl14-cKO retinas at 2 hours after EdU injection. Scale bar: 50μm. **(D)** Ki67 staining of peripheral areas of control and Mettl14-cKO retinas at P10. D2, D4, D6 and D8 are higher magnification views of the highlighted regions in D1, D3, D5 and D7. Scale bar: 200μm. **(E)** The number of Caspase-3 positive cells in control and Mettl14-cKO retinas from P3 to P21. Mean ± SD (n=3-9; *P<0.05, **P<0.01, ****p< 0.0001; Two-tailed Student’s T test). **(F)** The number of TUNEL positive cells in control and Mettl14-cKO retinas at P14, P21 and P56. Mean ± SD (n=3-9; ****p< 0.0001; Two-tailed Student’s T test). **(G)** The schematic showing the developmental defects and neuronal cell death observed in Mettl14-cKO retinas compared to controls.

To further confirm the role of *Mettl14* on cell survival in differentiated retina, we examined the level of cell death in Mettl14-cKO and control retinas (Figure 2E-F and Figure S3) at P14, P21 and P56 using caspase3 staining (detecting apoptosis) and TUNEL assay (marking end stage cell death). At P14, retinal differentiation is finished. The number of caspase3 positive cells was significantly higher at P14 and P21 in Mettl14-cKO retinas compared to controls. The TUNEL signals were significantly increased at P21 in Mettl14-cKO retinas. We also quantified the number of different retinal cell types at P14, P21 and P56 (Figure 1F & S4). The number of early-born neurons (born before birth), including retinal ganglion cells (RGCs) and horizontal cells (HCs), stayed comparable between Mettl14-cKO and control retinas at P14, but significantly decreased in Mettl14-cKO retinas starting from P21. The Mettl14-cKO retinas contained significantly less amacrine (ACs, born at both embryonic and neonatal stages) and late-born bipolar cells (BPs, born after birth) at all three stages (Figure 1F). The number of photoreceptor cells were also significantly reduced in Mettl14-cKO compared to control as indicated by the reduced thickness of ONL (Figure 1E). Notably, the number of Sox9 positive Müller glial cells (MGs) was consistently higher in Mettl14-cKO compared to control across all three stages (Figure 1F). Therefore, Mettl14 depletion significantly affected the survival of retinal neurons but not glial cells in differentiated retinas.

Overall, we detected delayed retinal development and increased neuronal cell death in differentiated retina when Mettl14-mediated m^6^A modification was depleted (Figure 2G). Interestingly, Mettl14 appeared to be only indispensable for postnatal retinal development. To further support this notion, we deleted the *Mettl14* gene in RPCs starting from P0 by in vivo electroporation. Electroporation primarily targets mitotic RPCs because nuclear envelope breakdown is necessary for plasmids to enter nuclei (Matsuda and Cepko 2007, 2004). We co-electroporated the *CAG-Cre* and *CAG-LoxP-stop-LoxP-GFP* plasmids into P0 *Mettl14^f/f^* or WT (littermates) mouse retinas in vivo (Figure S5). The *Mettl14^f/f^* retinal cells expressing Cre contained *Mettl14* deletion and were highlighted by GFP expression. The WT retinas received the same plasmid mix were served as controls. At P7, we observed significantly number of Ki67+ cells in the electroporated part of the retina in the presence of Cre, but not in control retinas (Figure S5 B-C). This phenocopies Mettl14-cKO retinas at P7 and further confirms that *Mettl14* is required for retinal development at postnatal stages.

### Mettll4-depleted late-born RPCs have extended cell cycle duration

While Mettl14 depletion affects both retinal development and neuronal survival, this manuscript primarily focuses on dissecting the role of Mettl14-mediated m^6^A modification during retinal development. To further investigate how *Mettl14* depletion affects retinal development at postnatal stages, we first traced the excessive RPCs found at P7 in Mettl14-cKO retinas using EdU pulse-chase assays. As mouse pups often die after intraperitoneal injection of EdU, we injected EdU via the intravitreal route. All the mouse pups survived after intravitreal injection of EdU and we did not detect significant toxicity effects in controls. We injected EdU into Mettl14-cKO and control mice at P7 and harvested the retinas at 2 hours post EdU injection (P7), P8 (24 hours post EdU injection), P9 (48 hours after EdU injection) and P10 (72 hours after EdU injection) (Figure 3A-D).

**Figure 3.**
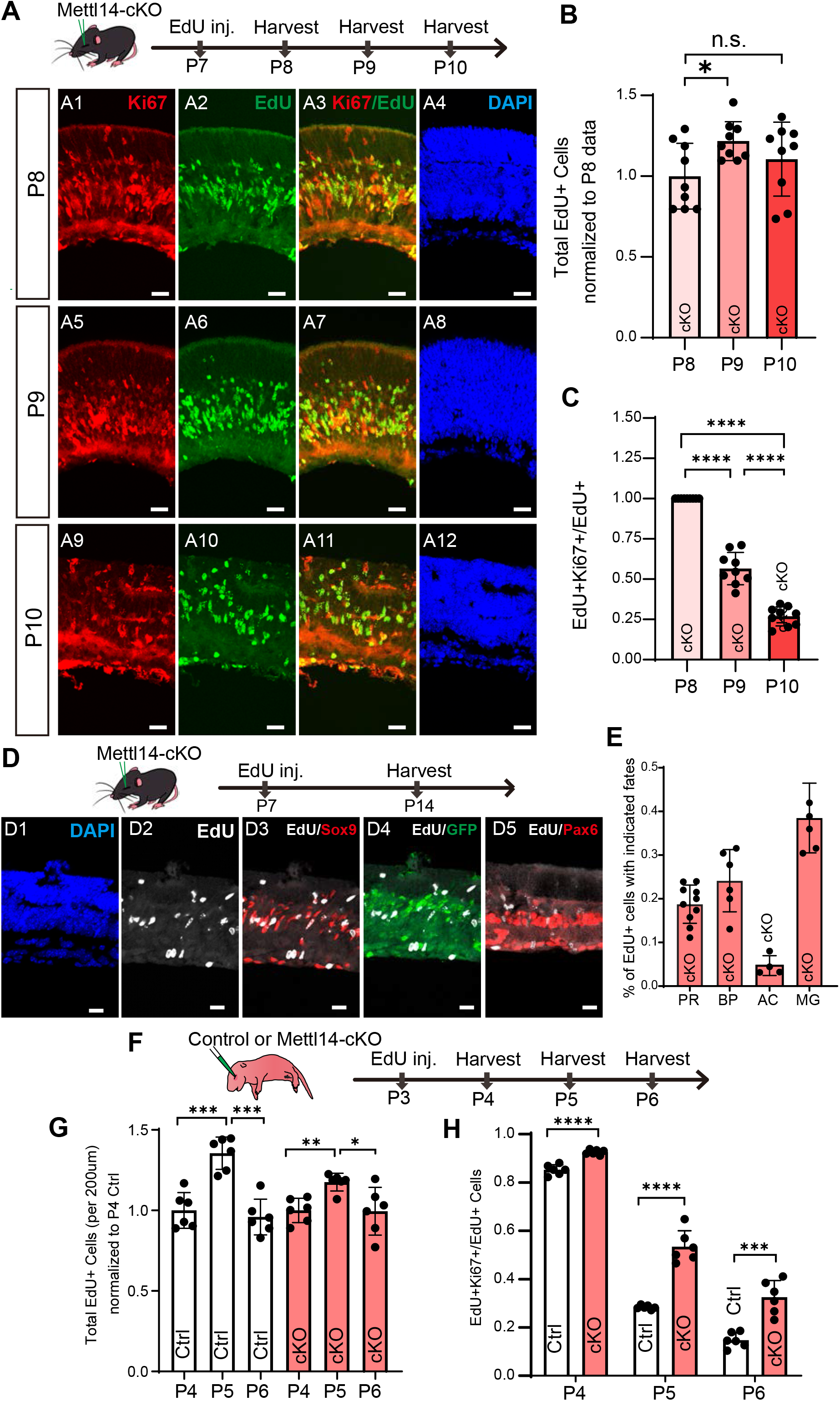
Mettl14-depleted late-born RPCs have extended cell cycle duration. **(A)** The EdU pulse-chase experiment in P7 Mettl14-cKO mice in vivo. The experimental design is shown on top. Scale bar: 20μm. **(B)** The number of EdU+ cells at P8, P9 and P10 in Mettl14-cKO retinas after EdU injection at P7. Data were normalized to values at P8. Mean ± SD (n=6, ** P< 0.01; Two tailed Student’s T test). **(C)** The percentage of cells remained in cell cycle (Ki67+EdU+/EdU+ cells) at P8, P9 and P10 in Mettl14-cKO retinas after EdU injection at P7. Data were normalized to the values at P8. Mean ± SD (n=6, ****P< 0.0001; Two tailed Student’s T test). **(D)** The fates of EdU+ cells in Mettl14-cKO at P14 after EdU injection at P7. The experimental design is shown on top. Scale bar: 20μm. GFP expression is driven by the Chx10 promoter in transgenic mice (*Chx10-EGFP/Cre*). GFP+Sox9-negative cells were counted as bipolar cells. **(F)** The percentage of EdU+ cells that adopted indicated cell fates at P14. BP: bipolar cells; AC: amacrine cells; MG: Müller glial cells. Mean ± SD (n>=4). **(F)** The EdU pulse-chase experiment in P3 control and Mettl14-cKO mice in vivo. **(G)** The number of EdU+ cells at P4, P5 and P6 in Mettl14-cKO retinas after EdU injection at P3. Data were normalized to values at P4 in controls. Mean ± SD (n=6, * P<0.05; ** P< 0.01; *** P<0.001; Two tailed Student’s T test). **(H)** The percentage of cells remained in cell cycle (Ki67+EdU+/EdU+ cells) at P4, P5 and P6 in control and Mettl14-cKO retinas after EdU injection at P3. Mean ± SD (n=6, *** P<0.001; ****P< 0.0001; Two tailed Student’s T test).

Retinal development progresses from the central to peripheral regions (Venters et al. 2015). Cell differentiation has finished in the central and mid-peripheral regions of the retina by P7 in wildtype mice. We did not observe any EdU+ cells in these regions in control mice at or after P7 (Figure 2C3, 2 hours post EdU injection). In Mettl14-cKO retinas, a significant number of EdU+ cells were found in the central and mid-peripheral regions of the retina at P7 (Figure 2C7, 2 hours post EdU injection). At 72 hours post EdU injection (P10), the number of EdU+ cells in Mettl14-cKO was not significantly increased compared to that at 24 hours post EdU injection (P8), suggesting that EdU+ P7 RPCs did not undergo excessive division in Mettl14-cKO retinas (Figure 3B). The number of the EdU+ cells remained Ki67 positive gradually decreased at P9 and P10 compared to P8, indicating that P7 Mettl14-cKO RPCs slowly exited cell cycle (Figure 3D). These results suggest that P7 Mettl14-cKO RPCs have an extended cell cycle duration.

To further confirm this notion, we traced RPCs at P3 in both control and Mettl14-cKO retinas using EdU pulse-chase assays (Figure 3F-G and S6). We injected EdU into Mettl14-cKO and control mice at P3 and harvested the retinas at P4 (24 hours post EdU injection), P5 (48 hours post EdU injection) and P6 (72 hours after EdU injection). In Mettl14-cKO retinas, the total number of EdU+ cells did not increase as dramatic as in controls at P5 compared to P4 (Figure 3G), suggesting that the Mettl14-cKO RPCs are less proliferative. Please note that the decreased number of EdU+ cells at P6 compared to P5 in control retinas may be due to the naturally occurring apoptosis peak at P5 (Figure 2E). Like P7 Mettl14-cKO RPCs (Figure 3A-C), EdU labeled P3 Mettl14-cKO RPCs exited the cell cycle slower than control RPCs (Figure 3H). There was significantly increased number of EdU+ cells staying Ki67+ in Mettl14-cKO retinas compared to controls at P5 and P6 (Figure 3H). Therefore, P3 Mettl14-cKO RPCs also proliferate less and exit the cell cycle slower than control RPCs. These results further support that Mettl14-cKO RPCs have extended cell cycle duration.

In addition, we investigated whether extended cell cycle duration would affect the fates of RPCs in Mettl14-cKO retinas. We labeled the excessive P7 RPCs in Mettl14-cKO retinas by EdU and traced their fates at P14 when we can confidently assess cell fates by marker expression (Figure 3D-E). We found that they had mainly differentiated into late-born bipolar cells (~23%) and Müller glial cells (~38%) (Figure 3E). Compared to the fates of wild type P2 and P5 RPCs in the published literature, the P7 Mettl14-cKO RPCs cells are biased towards adopting late-born cell fates (Figure S7).

Overall, we show that late-born RPCs have extended cell cycle and favor late-born bipolar and Müller glial fates in the absence of *Mettl14.*

### m^6^A directly regulates cell cycle-related genes in the developing retina

To investigate the molecular mechanisms underlying *Mettl14*-depletion-induced late-born RPC cell cycle progression delay, we profiled m^6^A modified mRNAs in the developing retina at P7 and searched for mRNAs whose m^6^A levels were eliminated with *Mettl14* deletion. Methylated RNA immunoprecipitation sequencing (MeRIP-seq or m^6^A-seq), which uses an antibody against m^6^A to immunoprecipitate m^6^A-enriched mRNAs, allows for mapping of m^6^A modified mRNAs in a genome-wide manner (Delatte et al. 2016). We examined P7 control and Mettl14-cKO retinas using m^6^A-seq in combination with regular RNA-seq.

The m^6^A modification was found to be enriched at the consensus RRACH motif (R=G or A; H=G, A or C) in both control and Mettl14-cKO retinas at P7, which is consistent with studies in other systems (Figure 4A) (Chen et al. 2015; Zhang et al. 2019). The overall landscape of m^6^A was not significantly changed in Mettl14-cKO, with m^6^A modified sites primarily locating at CDS (coding DNA sequence), stop codon and 3’UTR (3’ untranslated region) (Figure 4B). In control P7 retinas, mRNAs of approximately 79% of expressed genes contained significant levels of m^6^A modification (Figure 4C). These m^6^A-modified mRNAs are highly enriched for involvement in pathways such as neurogenesis and stem cell differentiation (Figure 4D). Compared to controls, a total of 1269 genes were differentially expressed in Mettl14-cKO retinas. Among the 482 downregulated genes, approximately 12% of the genes showed significantly reduced m^6^A levels. Gene Ontology analyses showed that these downregulated genes were highly involved in circadian regulation of gene expression, blood vessel remodeling and visual system development (Figure 4G). However, the reduction of m^6^A levels showed no correlation with changes of mRNA levels (data now shown). Among the 787 significantly upregulated genes, the mRNAs of 101 genes were m^6^A modified and showed significantly reduced m^6^A enrichment in Mettl14-cKO retinas. For these genes, the reduction of m^6^A levels was negatively correlated with changes of mRNA levels, which is consistent with the major role of m^6^A modification in mediating mRNA decay (Figure 4E). We focused on these genes, hypothesizing that they are likely to be direct targets of m^6^A modification. Gene Ontology analyses revealed that these genes were highly involved in cell cycle related pathways, including cell division, regulation of cell cycle process, and cellular response to DNA damage stimulus (Figure 4F). We confirmed the increased mRNA levels of top candidate genes in Mettl14-cKO retinas using quantitative PCR (Figure 4H). Whereas some of these genes are involved in promoting cell cycle progression, many are known to slow down cell cycle, such as *Brca2, Brip1, Foxm1, Ttk* and *Birc5* (Figure 4H) (Gudmundsdottir and Ashworth 2006; Cantor and Guillemette 2011; Zona et al. 2014; Wei et al. 2005; Lens et al. 2003). The upregulation of these genes collectively led to extended cell cycle, as we observed in Mettl14-cKO retinas. Thus, *Mettl14* deletion-mediated m^6^A depletion resulted in sustained high levels of mRNAs that can slow down cell cycle, which induced late-born RPC cell cycle extension.

**Figure 4.**
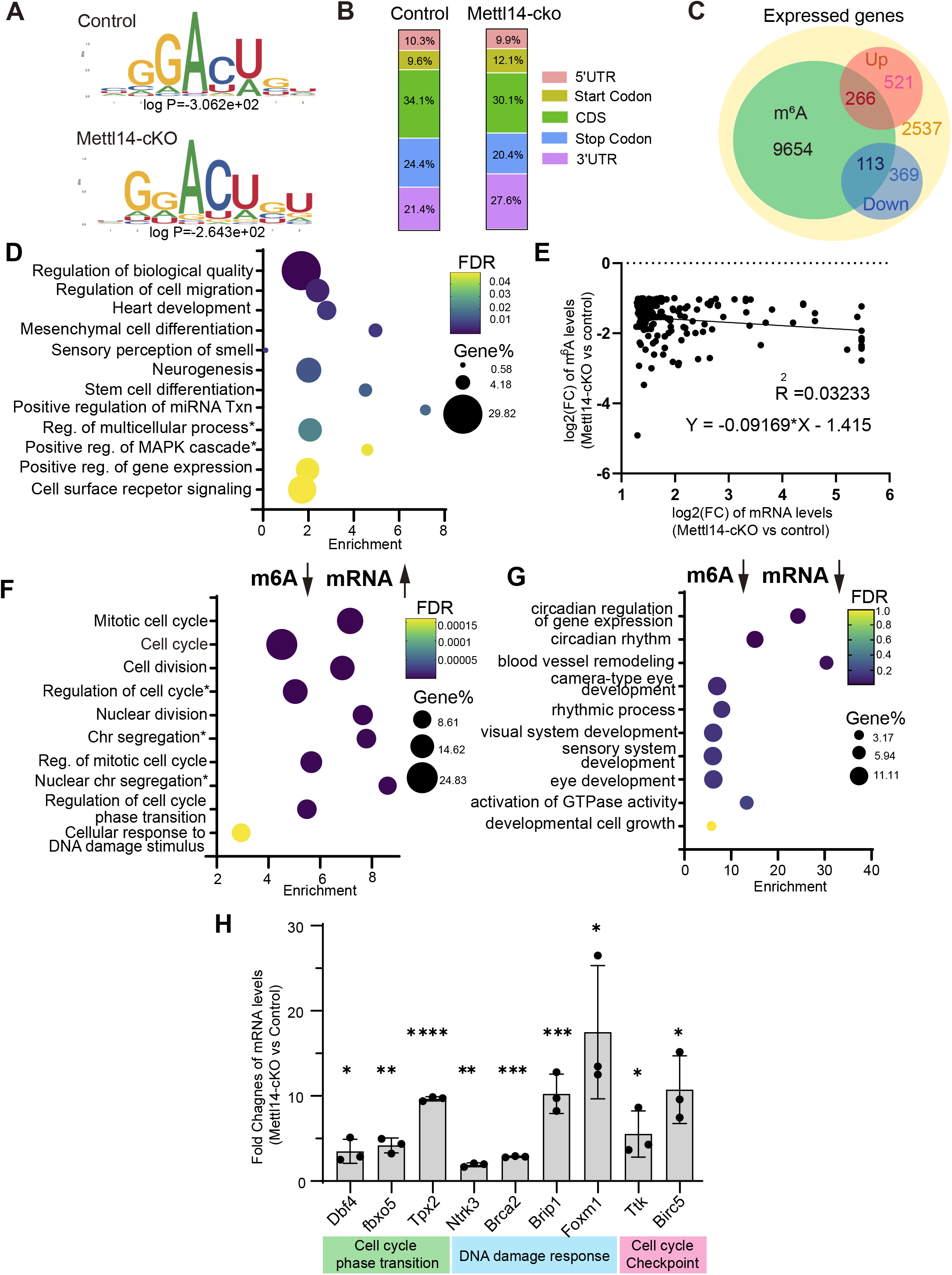
m^6^A-seq reveals that m^6^A directly regulates cell cycle related genes in the developing retina. **(A)** The consensus m^6^A motifs in control and Mettl14-cKO retinas at P7. **(B)** The m^6^A peak localization in control and Mettl14-cKO retinas at P7. **(C)** Yellow circle: the number of expressed genes in control retinas at P7. Green circle: the number of genes with m^6^A peaks in control retinas at P7. Red circle: the number of genes that are up-regulated in Mettl14-cKO retinas at P7 compared to controls. Blue circle: the number of genes that are down-regulated in Mettl14-cKO at P7. **(D)** The GO analysis of genes that are modified by m^6^A. **(E)** The linear regression analysis shows the correlation between m^6^A level decrease and mRNA level increase in P7 Mettl14-cKO retinas (P<0.05). **(F)** The GO analysis of genes with increased mRNA levels and decreased m^6^A levels in Mettl14-cKO retinas compared to controls at P7. **(G)** The GO analysis of genes with decreased mRNA levels and decreased m^6^A levels in Mettl14-cKO retinas compared to controls at P7. **(H)** QPCR verification of top candidate genes with increased mRNA levels and decreased m^6^A levels in Mettl14-cKO at P7. The known roles of the candidate genes are indicated in the boxes below the chart. All values were normalized to *Gapdh* levels. Data were also normalized to controls. Mean ± SD (n=3, * P< 0.05; ** P< 0.01; ***P< 0.001; ****p< 0.0001; Two tailed Student’s T test).

### Single cell RNA-seq reveals molecular events downstream of m^6^A modification in the developing retina

We performed single cell transcriptomic analyses (scRNA-seq via 10x Genomics) to further reveal *Mettl14* depletion-induced changes in late-born RPCs and different retinal cell types. We recovered 10451 cells from four P7 control retinas and 13116 cells from four Mettl14-cKO retinas. Based on well-established markers for retinal cell types (Clark et al. 2019), we assigned known cell identities to the 25 clusters grouped by an unbiased clustering method (Seurat 4.1) (Figure S8). The 11 RPC-derived cell types were selected for downstream analyses (Figure 5A-C). In addition to the seven major retinal cell types (rod, cone, Müller glia, bipolar, amacrine, horizontal and ganglion cells), we detected cell clusters representing RPCs and precursors of rod, Müller glia, and bipolar cells in P7 retinas. We focused on analyzing the RPC and precursor cell clusters to elucidate the mechanisms underlying m^6^A regulation.

**Figure 5.**
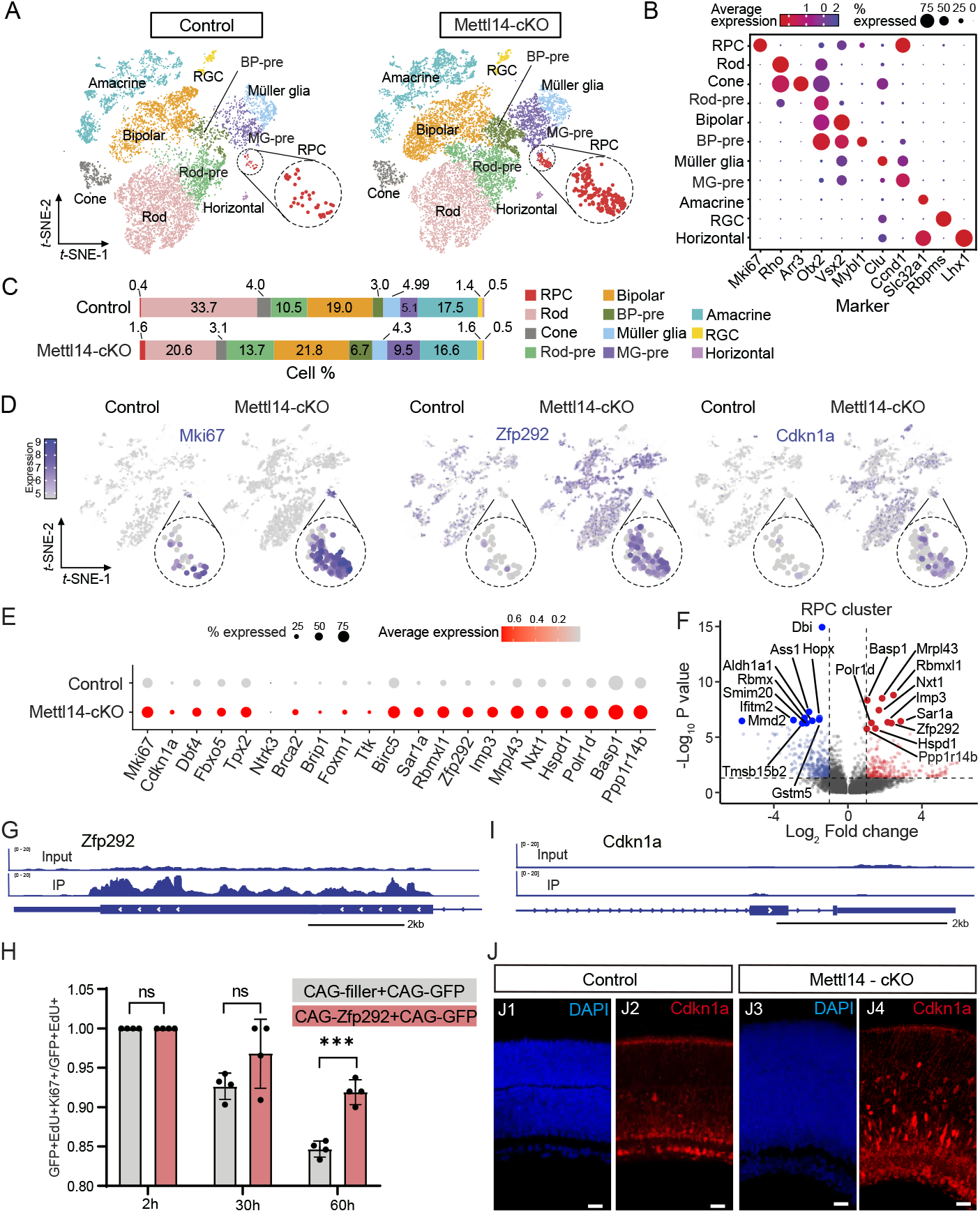
Single cell RNA-seq reveals molecular events downstream of m^6^A modification in the developing retina. **(A)** T-SNE maps of retinal cell types captured by scRNA-seq in control (left) and Mettl14-cKO (right) retinas at P7. The cell clusters of RPCs are circled and highlighted as inserts. **B)** Dot plot showing the markers used for identifying retinal cell types. Dot size: the proportion of cells expressing indicated marker genes. Dot color: the average expression levels. RPC: retinal progenitor cell; Rod-pre: rod precursor; BP-pre: bipolar precursor; MG-pre: Müller glia precursor; RGC: retinal ganglion cells. **(C)** The proportion of each cell cluster in control (Top) and Mettl14-cKO (Bottom) retinas at P7. **(D)** Feature plots showing the expression of *Mki67* (left), *Zfp292* (middle) and *Cdkn1a* (P21) (right) in control and Mettl14-cKO retinas at P7. The cell cluster of RPCs is circled and highlighted as inserts. **(E)** Dot plot showing the expression of cell cycle-related genes (qPCR verified in Figure 3G), and top 10 upregulated genes in RPC clusters (Mettl14-cKO vs Control). **(F)** Volcano plot showing differential expressed genes (DEGs) between Mettl14-cKO and control RPCs at P7. DEGs that are up-or down-regulated in Mettl14-cKO are illustrated in light red or blue, respectively. The cutoff for DEGs is p < 0.05 (Wilcoxon rank sum test) and |log2 fold change| >1 (shown as dash lines). The 10 most significantly upregulated or downregulated DEGs are highlighted in red or blue respectively. **(G)** m^6^A peaks on the *Zfp292* gene in wildtype retinas at P7. **(H)** Two-hour, 30-hour and 60-hour EdU pulse-chase of P0 RPCs overexpressing *Zfp292* in retina explants. Mean ± SD (n=4, ***P< 0.001; Two tailed Student’s T test). *CAG-filler*: filler plasmid DNA to ensure the equivalent plasmid concentrations. **(I)** m^6^A peaks on the *Cdkn1a* gene in wildtype retinas at P7. **(J)** P21 (encoded by the *Cdkn1a* gene) staining in control and Mettl14-cKO retinas at P7. Scale bar: 20μm.

In Mettl14-cKO retinas, there were increased number of *Mki67*+ (the gene encoding Ki67) RPCs compared to controls (Figure 5C). Some *Mki67+* cells were also found in Müller glia and bipolar cell precursor clusters in Mettl14-cKO but not in controls (Figure 5D). This increased number of *Mki67*+ cells is consistent with the observations described in Figure 1 – 3. In addition, the percentage of retinal cells differentiating towards bipolar and Müller glial fates (BP+BP-pre, MG+MG-pre) was increased in Mettl14-cKO retinas compared to controls, whereas the percentage of cells differentiating towards rod fate (Rod+Rod-pre) was decreased. This matches our finding that P7 Mettl14-cKO RPCs favor bipolar and Müller glial fates (Figure 3D&E). Notably, the proportions of the precursor cells for rod, bipolar, and Müller glial cells were all increased in Mettl14-cKO retinas, suggesting a development delay. The proportions of the cell types derived from early-RPCs, such as ganglion cells, cones, and horizontal cells, were not changed in Mettl14-cKO retinas compared to control retinas (Figure 5C).

We analyzed the genes that were differentially expressed in the RPC cluster in control and Mettl14-cKO retinas. The cell cycle-related genes revealed by m^6^A-seq and bulk RNA-seq (Figure 4H) showed increased expression in Mettl14-cKO RPCs compared to control RPCs (Figure 5E). The top 10 genes that were significantly upregulated in Mettl14-cKO RPCs were highlighted in Figure 5E and 5F. Most of these genes showed reduced m^6^A levels based on m^6^A-seq (Figure S9A). Some of these genes have been implicated in regulating the cell cycle in other systems(Zhou et al. 2018; Wang et al. 2019; Liang et al. 2014; Bhargava et al. 2018; Gong et al. 2021), but their roles in RPC cell cycle progression have not been well studied. The top candidate was *Zfp292,* which encodes the transcription factor Zinc Finger Protein 292 (also known as Znf292). It was highly modified by m^6^A in control retinas, and its m6A level was significantly reduced in Mettl14- cKO retinas (Figure 5G & S9A). Several studies have identified *Zfp292* as a tumor suppressor gene (Takeda et al. 2015; Jh et al. 2016; Gong et al. 2021), but its function in the retina or the CNS is unknown. We over-expressed *Zfp292* in wildtype late-born RPCs and examined whether sustained expression of *Zfp292* can inhibit the cell cycle progression of late-born RPCs. The plasmid over-expressing *Zfp292* (*CAG-Zfp292*) was delivered into P0 mouse retinas *ex vivo* via electroporation. The *CAG-GFP* plasmids were co-electroporated with the *CAG-Zfp292* plasmids to label the cells over-expressing *Zfp292*. The efficiency of co-electroporating two types of plasmids into the same cells exceeds 90% (Matsuda and Cepko 2004, 2007). Twenty-four hours after retinas were electroporated with the control or *CAG-Zfp292* plasmids, we applied EdU to retinas for 1 hour and harvested them 2, 30, or 60 hours later to trace the cell cycle progression of late-born RPCs (Figure 5H &S10). As development proceeded, the GFP+EdU+ cells gradually exited the cell cycle and became Ki67 negative. The percentage of GFP+EdU+ cells that were Ki67+ continuously decreased at 30 and 60 hours after the EdU pulse in controls, while the percentage remained high in retinas over-expressing *Zfp292.* This demonstrated that the RPCs over-expressing *Zfp292* failed to exit the cell cycle normally as development proceeded. The total number of GFP+EdU+ cells was not significantly higher when *Zfp292* was over-expressed compared to controls, suggesting that *Zfp292* did not promote RPC proliferation. Based on these results, we concluded that *Zfp292* over-expression can significantly inhibit late-born RPC cell cycle progression. The increased levels of *Zfp292* mRNAs in Mettl14-cKO retinas thus can contribute to RPC cell cycle extension.

We also analyzed differentially expressed genes in clusters classified as precursors for rods, bipolar cells, or Müller glial cells. Interestingly, *Cdkn1a,* which encodes the P21^Cip1^ protein and is well-known for its role in inhibiting cell cycle progression (Karimian et al. 2016), was significantly upregulated in all three precursor clusters in Mettl14-cKO retinas (Figure 5D&S9B-D). Notably, the *Cdkn1a* mRNAs were unlikely to be directly regulated by m^6^A because m^6^A-seq did not detect m^6^A enrichment on *Cdkn1a* mRNAs (Figure 5I). The increased level of *Cdkn1a* is likely to be an indirect event induced by m^6^A depletion. Interestingly, while not statistically significant in scRNA-seq data, *Cdkn1a* mRNA levels appeared to be higher in the RPC cluster in Mettl14-cKO retinas compared to control retinas (Figure 5D). We confirmed the increased levels of *Cdkn1a* mRNA and protein in Mettl14-cKO retinas at P7 using qPCR, *in situ* RNA hybridization, and immunofluorescent staining (Figure 5J & S9E-F). It is likely that this m^6^A depletion induced P21^Cip1^ upregulation also contributes to RPC cell cycle extension in Mettl14-cKO retinas.

In addition, we compared the P7 Mettl14-cKO RPCs with wildtype P0, P2 and P8 RPCs in published scRNA-seq datasets (Figure S11A) (Clark et al. 2019). The P7 Mettl14-cKO RPCs intermingled with wildtype RPCs as shown in the UMAP (Uniform Manifold Approximation and Projection) plot, suggesting that they are not dramatically different from wildtype RPCs at the transcript levels (Figure S11B-E). The cell-cycle scoring and regression analyses (Seurat 4. 0. 6) revealed that the percentage of P7 Mettl14 RPCs in the G2/M phases of the cell cycle was higher than that in P7 control RPCs (Figure S11F). This indicates that Mettl14-cKO RPCs may be slowed down in the G2/M phases. Notably, P7 Mettl14-cKO RPCs differentially expressed genes that are involved in the cellular senescence pathway (Figure S11G). Given that P21^Cip1^ is a well-established senescence marker (Kumari and Jat 2021), this result further suggests that P21^Cip1^-related pathways may play important roles downstream of m^6^A depletion.

Taken together, the scRNA-seq analyses further confirmed our findings and revealed *Zfp292* and *Cdkn1a* to be important contributors of m^6^A depletion-induced late-born RPC cell cycle extension.

## Discussion

We investigated mechanisms countering the RPC cell cycle lengthening during retinal development. We showed that Mettl14-mediated m^6^A modification is critical for ensuring the proper cell cycle progression of late-born RPCs. Depletion of m^6^A modification by conditionally deleting the *Mettl14* gene in the developing retina led to cell cycle extension. m^6^A-seq and scRNA-seq revealed that mRNAs involved in slowing down cell cycle were highly enriched for m^6^A, and their levels were significantly higher in the absence of *Mettl14* in late-born RPCs. In addition, we identified *Zfp292* to be a novel target of m^6^A in the developing retina and suggested that P21^Cip1^-related pathways were involved in slowing down the cell cycle upon m^6^A depletion. Notably, m^6^A modification appeared to be dispensable for RPCs that were born before birth, because no significant defects were detected in the retina prior to or at birth. Based on these results, we propose a model for how late-born RPCs counter the cell cycle lengthening during retinal development. Under wildtype conditions, the mRNAs slowing down cell cycle progression, such as *Zfp292,* are modified by m^6^A, which decreases their mRNA levels possibly through the mRNA decay pathway and can avoid uncontrolled cell cycle lengthening. In the absence of m^6^A, the levels of these mRNAs are higher, resulting in cell cycle extension.

While it has not been studied in the developing retina, the function of m^6^A modification on neural development has been reported. During cortical neurogenesis, depletion of m^6^A via deleting different components of the m^6^A machinery resulted in two major outcomes. First, a prolonged cell cycle of embryonic neural progenitor cells and extended neurogenesis were found in *Mettl14* or *Fmr1* (encodes a protein reading m^6^A and mediating mRNA nuclear export) conditional knock out mice (Yoon et al. 2017; Edens et al. 2019). Second, deletion of *Mettl14* or *Ythdf2* (encoding a protein reading m^6^A and primarily mediating mRNA decay) resulted in reduced proliferation and premature differentiation of neural progenitor cells (Wang et al. 2018; Li et al. 2018). During cerebellar development, m^6^A modification downregulated apoptosis-related mRNAs and ensured the survival of newly generated cerebellar granule cells (Wang et al. 2018a). We observed extended cell cycle duration when Mettl14-medidated m^6^A was depleted in RPCs. We further showed that this m^6^A regulation is required in late-born RPCs with long cell cycle duration, but appeared to be dispensable for early-born RPCs with short cell cycle duration. While intensively studied, our understanding of the heterogeneity and competence of RPCs is still limited. Recent single cell transcriptomic analyses revealed that there are clearly two distinct populations of RPCs, which are the ones born before or after E16-E18 (Clark et al. 2019). Our work shows that RPCs born after birth (late-born RPCs) are sensitive to m^6^A regulation, and further highlights the heterogeneity among RPCs.

This study also identified a novel target of m^6^A in the developing retina, *Zfp292,* which can significantly inhibit the cell cycle progression of late-born RPCs. We further showed that *Cdkn1a* mRNAs and proteins (P21^Cip1^) were significantly upregulated in the retina upon m^6^A depletion. P21^Cip1^ is well-known for its role in inhibiting cell cycle progression. Interestingly, the upregulation of P21^Cip1^ was not a direct effect of m^6^A depletion, because *Cdkn1a* mRNAs were not highly enriched for m^6^A modification in wild type retinas. This increased expression of P21^Cip1^ is also not directly downstream of *Zfp292* in that over-expression of *Zfp292* did not lead to upregulation of *Cdkn1a* in wild type retinas (data not shown). One possibility is that P21^Cip1^ expression is induced or accompanied by the upregulation of genes involved in the DNA damage response pathways, which were found to be modified by m^6^A and their levels were significantly increased in Mettl14-cKO retinas (Figure 4F&H). Most of these genes are known to slow down cell cycle during DNA damage repair. We did not detect obvious DNA damage in Mettl14-cKO retinas at neonatal stages (data not shown). The upregulation of these genes is likely a direct effect of m^6^A depletion. Thus, it is possible that late-born RPCs express genes involved in DNA damage response pathways to prime themselves for potential DNA damage during proliferation. m^6^A could be one of the mechanisms that is used by late-born RPCs to balance the levels of these genes and ensure normal cell cycle progression.

A caveat of this study is that we were not able to directly confirm whether Mettl14- mediated m^6^A modification functions through the mRNA decay pathway in the retina. We tried to measure the mRNA half life following published protocols. However, we noticed that most retinal cells died quickly when we placed the retina inside the Actinomycin solution (inhibiting transcription).

In addition, we observed significant neuronal cell death in mature Mettl14-cKO retinas. While this is not the focus of this manuscript, we performed m^6^A-seq on P30 control and Mettl14-cKO retinas (Figure S12). The mRNAs of genes involving in synapse organization and assembly were highly enriched for m^6^A modification (Figure S12C). The levels of mRNAs involving in regulating cell morphogenesis were significantly increased in Mettl14-cKO retinas (Figure S12D). The levels of mRNAs involving in histone modification was significantly downregulated in Mettl14-cKO (Figure S12E). It would be important to elucidate how these m^6^A modified mRNAs contribute to neuronal cell death in mature retinas in future studies.

In summary, we studied how RPCs counter cell cycle lengthening and ensure proper cell cycle progression. We showed that post-transcriptional m^6^A modification on mRNAs plays a key role in this process. In the future, it will be important to investigate how cell cycle duration is differentially regulated in early and late RPCs, how cell cycle lengthening is coordinated with the differentiation program, and how m^6^A regulation interacts with these programs to guarantee proper development. It will also be interesting to further investigate the function of *Zfp292* during retinal development and examine whether it can serve as a therapeutic target to control cell cycle progression under disease conditions.

## Experimental procedures

### Animals

The Mettl14^f/f^ mouse line was a gift from Dr. Chuan He at the University of Chicago (Yoon et al. 2017). The Chx10-EGFP/Cre mice were purchased from the Jackson Lab (Stock# 005105). Wild-type mouse neonates were obtained from timed pregnant CD1 mice (Charles River Laboratories, #022). All animal studies were approved by the Administrative Panel on Laboratory Animal Care (APLAC) at Stanford University.

### Plasmid Construction

The *CAG-GFP, CAG-LoxP-Stop-LoxP-GFP, CAG-Cre* and *CAG-filler* plasmids were from Wang et al. 2014. To generate the *CAG-Zfp292* plasmid, we split the mouse *Zfp292* ORF into four overlapping DNA fragments (each about ~2kb), which were then synthesized by IDT. We cloned these fragments into the *CAG-GFP* backbone by Gibson Assembly (NEB, Cat.No. E2611). Primers are listed in Table S1&2.

### Western blot

Mouse retinas were freshly dissected out, transferred to RIPA buffer (Radioimmunoprecipitation assay buffer, Abcam #ab156034), and heated at 95°C for 10 mins. Protein concentrations were measured by BCA protein kit (Pierce™ BCA Protein Assay Kit #23228). Equal amounts of proteins were loaded onto a 4%-20% SDS-PAGE gel (BIO-RAD, #4561094) and run at 130V and 30A for 1.5 hours. The gel was transferred to a 0.2mm PVDF membrane (BIO-RAD, #1704156) using a semi-dry transfer system (BIO-RAD, 1704150EDU). The membrane was blocked with 5% non-fat milk blocking solution and incubated with primary antibodies at 4°C overnight. Primary antibodies included rabbit anti-Mettl14 (Sigma-Aldrich, HPA038002 1:1000) and rabbit anti-GAPDH (Abcam, ab9485, 1:1000). The membrane was washed with 1X TBST (20mM Tris, 150mM NaCl, 0.1% Tween20) for 30 mins and incubated with secondary antibody conjugated to horseradish peroxidase (Amersham, NA934-100U, 1:5000) at room temperature for 2 hours. The membrane was washed again and incubated with the SuperSignal West Pico PLUS chemiluminescent substrate mix (Thermo Scientific, #34580). Chemiluminescent signals were detected using the Amersham ImageQuant 600 (Cytiva) imaging system.

### Quantitative real time PCR

Total RNA was isolated from retinal tissues using QIAGEN RNeasy kit (QIAGEN, 74104). The cDNA was generated using SuperScript™ III First-Strand Synthesis Kit (Invitrogen, 18080051). qPCR primers were listed in Table S1. The SYBR™ Green Master Mix (Fisher Scientific, A25742) was used for qPCR. The qPCR experiments were performed using a Quantstudio3 Real-Time PCR System (Applied Biosystems, A28567) with the following conditions: 50°C for 2 mins, 95°C for 10 mins, followed by 40 cycles of 95°C for 15s, 60°C for 1min.

### Ex vivo and in vivo plasmid electroporation into the retina

Ex vivo and in vivo retina electroporation was carried out as previously described (Matsuda and Cepko 2004, 2007; Wang et al. 2014; Emerson and Cepko 2011). For ex vivo electroporation, 5 pulses of 25V, 50ms each with 950ms intervals were applied to dissected retinas. For in vivo electroporation, 5 pulses of 80V, 50ms each with 950ms intervals were applied to neonatal mouse pups. All ex vivo and in vivo electroporation experiments were repeated with at least three biological replicates. Plasmids were electroporated with a concentration of 500ng/ul to 1ug/ul per plasmid.

### Histology and immunofluorescence

Mouse eyeballs were fixed in 4% paraformaldehyde (PFA) in 1×PBS (pH 7.4) for 2 hours at room temperature. Retinas were then dissected out and equilibrated at room temperature in a series of sucrose solutions (5% sucrose in 1× PBS, 5 min; 15% sucrose in 1× PBS, 15 min; 30% sucrose in 1× PBS, 1 h; 1:1 mixed solution of OCT and 30% sucrose in PBS, 4°C, overnight), embedded in OCT and stored at −80°C. A Leica CM3050S cryostat (Leica Microsystems) was used to prepare 20 μm cryosections. Retinal cryosections were washed in 1× PBS briefly, incubated in 0.2% Triton, 1× PBS for 20 min, and blocked for 30 min in blocking solution (0.1% Triton, 1% BSA and 10% donkey serum in 1x PBS, Jackson Immuno Research Laboratories). Slides were incubated with primary antibodies diluted in blocking solution in a humidified chamber at 4°C overnight. After washing in 0.1% Triton 1× PBS for three times, slides were incubated with secondary antibodies and DAPI (Sigma-Aldrich; D9542) for 1 hour, then washed for three times with 0.1% Triton, 1× PBS and mounted in Fluoromount-G (Southern Biotechnology Associates). The following primary antibodies were used: chicken anti-GFP (Abcam, ab13970, 1:1000), rabbit anti-Sox9 (Abcam, ab185966, 1:500), rabbit anti- Pax6 (Thermo Fisher Scientific, 42-6600, 1:500), mouse anti-Rhodopsin (Abcam, ab5417, 1:200), mouse anti-Bassoon (Abcam, ab82958, 1:200), mouse anti-Ki67 (BD Biosciences, 550609, 1:200), rabbit anti-P21 (1:500, Abcam, ab188224), Sheep anti- Chx10 (Exalpha Biologicals, X1179P, 1:100), Gunine Pig anti-RBPMS (PhosphoSolutions, 1832-RBPMS, 1:500), mouse anti-Calbindin (Sigma-Aldrich, C9848, 1:500), mouse anti-Rhodopsin (Abcam, ab5417, 1:200), Rabbit anti-GFAP (Fisher Scientific, NC9604017, 1:200) and rabbit anti-Mettl14 (Sigma-Aldrich, HPA038002, 1:500).

### TUNEL assay

TUNEL Assay was performed using the Click-iT™ Plus TUNEL Assay for In Situ Apoptosis Detection, Alexa Fluor™ 647 dye kit (Invitrogen™ C10619).

### EdU pulse-chase assay

EdU (5μM) was injected intravitreally into mice pups at P3 and P7 using the femtoJet system. EdU signals were detected on retinal cryosections using the EdU Cell Proliferation Kit (Invitrogen, C10340).

### m^6^A dot-blot assay

Total RNA samples were denatured at 95°C, spotted onto Hybond-N+ membrane (Catalog No. GERPN1510B; GE Healthcare) and crosslinked using a Stratalinker 2400 UV crosslinker. Membranes were blocked with 5% milk blocking solution for 1 hour, and then incubated with anti-m^6^A antibody (Synaptic Systems, 202003) for 2 hours at room temperature. Membranes were washed and incubated with HRP-conjugated secondary antibodies for 1 hour at room temperature. For signal detection, membrane was incubated with ECL Western Blotting Substrate for 5 min in darkness at room temperature and then imaged using the Amersham ImageQuant 600 (Cytiva) imaging system.

### RNA in situ hybridization

RNA in situ hybridization was performed using the Yn-situ method (Wu et al. 2022). Briefly, the retinal cryo-sections were fixed with 4% PFA for 1 hour, followed by EDC (1- Ethyl-3-(3-dimethylaminopropyl) carbodiimide) treatment for 1 hour at room temperature. For hybridization, sections were digested with proteinase K (1:200 dilution) at 40°C for 30 mins and blocked with hybridization buffer for 5 min. Probe mixtures were added onto the sections, which were incubated at 40°C for overnight. Slides were washed for three times with wash buffer (30% formamide, 5× SSC, 9 mM citric acid (pH 6.0), 0.1% Tween 20, 50 μg/mL heparin) and 5x SSCT (saline-sodium citrate, 0.1% Tween 20) and incubated in preamplifier (20 ng/μL) for 2 hours at 40°C. Signals were amplified by HCR hairpins (3uM) and imaged by the Zeiss confocal microscope (LSM880). Probes information can be found in Table S1.

### Imaging and analysis

All images of retinal sections were acquired by a Zeiss LSM880 inverted confocal microscope. Representative images were maximum projections of 5-10μm tissues and were quantified by Fiji software.

### m^6^A-seq (MeRIP-seq) and data analysis

Total RNA was extracted from retina tissues using RNeasy Mini Kit (QIAGEN, 74104). Approximately 30ug of total RNA per sample with RIN number >8.0 was used for m^6^A-seq. The poly(A) mRNAs were isolated by using the Dynabeads Oligo(dT) kit (ThermoFisher, #61002), fragmented into ~100bp-long oligonucleotides using divalent cations under elevated temperature, and incubated with m^6^A antibody (No. 202003, Synaptic Systems, Germany) in IP buffer (50 mM Tris-HCl, 750 mM NaCl and 0.5% Igepal CA-630) supplemented with BSA (0.5 μg μl-1) for 2 hours at 4°C. The mixture was then incubated with protein-A beads and eluted with elution buffer (1 × IP buffer and 6.7mM m6A). Eluted RNAs were precipitated by 75% ethanol. Eluted m6A-containing fragments (IP) and untreated input control fragments are converted to cDNA libraries and submitted for next generation sequencing (LC-BIO Bio-tech ltd, Hangzhou, China). The sequencing quality was verified by using FastQC. Bowtie was used to map the reads to the Mus musculus (Version: v96) genome with default parameters. Mapped reads of IP and input libraries were provided for R package exomePeak to identify m^6^A peaks adapted for visualization on the UCSC genome browser or IGV software. Motifs were identified and localized by using MEME scripts (Bailey et al. 2009). Called peaks were annotated by intersection with gene architecture using ChIPseeker (Yu et al. 2015). StringTie (Pertea et al. 2015) was used to perform expression analyses for all mRNAs from input libraries by calculating FPKM (total exon fragments/mapped reads(millions)×exon length(kB)). The differentially expressed mRNAs were selected with |log2 (fold change) |>1 and p value < 0.05 by R package edgeR. GO term analysis was performed by The Gene Ontology Resource (http://geneontology.org/).

### Single Cell RNA sequencing (scRNA-seq) and data analysis

Retinal tissues were dissociated as described previously (Wang et al. 2014) and loaded on a 10x Genomics Chromium controller to generate single cell Gel-Beads-in Emulsion (GEMs). Single-cell RNA-Seq libraries were prepared using the Chromium Gene Expression 3’ kit v3.1 (PN-1000269). The quality of the libraries was analyzed by a 2100 Bioanalyzer (Agilent) using a high-sensitivity DNA kit (Agilent, #5067-4626). The libraries were sequenced by Illumina HiSeq4000 platform (2 x 75 paired-end reads). Single cell RNA expression matrices were generated by aligning sequencing reads to a pre-build mouse reference (mm10-2020-A) using the Cell Ranger (v.6.1.1) pipeline and loaded into R (v.4.1) for downstream analyses. We applied SoupX (Young and Behjati 2020), DoubleDecon (DePasquale et al. 2019) and DoubletFinder (McGinnis et al. 2019) to remove ambient RNA and doublets. The filtered matrices were processed with Seurat (v.4.1) for quality control, clustering and visualization using standard workflows. Briefly, genes that were detected in fewer than 5 cells, and cells with fewer than 500 genes or more than 5% reads as mitochondrial were excluded from analyses. Control and Mettle14-cKO samples were normalized by SCTransform method (Hafemeister and Satija 2019) and integrated using 3000 variable genes (*PrepSCTIntegration, FindIntegrationAnchors and IntegrateData*). Clusters were generated based on the top 30 principal components (PCs) (*FindNeighbors* function and *FindClusters* function, resolution = 1) and visualized by t-distributed Stochastic Neighbor Embedding (tSNE). To identify the conserved markers of each cluster, we performed differential gene expression tests on each cluster versus all other clusters (*FindConservedMarkers*). Clusters were assigned to 15 major cell types by their conserved markers. Differentially expressed genes (DEGs) were identified using a Wilcoxon rank-sum test (*FindAllMarkers*) between control and Mettl14-cKO in each cell type. DEGs with ļlog2 (fold change)ļ > 1 and p value < 0.05 were selected as input for GO analysis described above. To query the P7 Mettl14- cKO RPCs on published reference datasets (Clark et al. 2019), the Unimodal UMAP projection function (*Mapquery*, Seurat 4.0.6) was used.

### Statistics

The sample size (n) was determined by using G*power software (two groups, the effect size of 0.5, α error at 0.05 and power set at 0.95). Personnel were blinded during experiments. GraphPad Prism v.9 (GraphPad) was used to perform statistical analysis. Student’s t-tests (two tailed) were used with the appropriate parameters. Statistical significance was defined as a P value <0.05.

### Data availability

Sequencing data have been uploaded to GEO database (GSE206013 and GSE206014).

## Acknowledgements

The members of the Wang and Ding laboratories provided valuable discussion and support for this project.

## Funding and additional information

Support was provided by NIH 1R01EY03258501 (to S.W. and L.L.) and P30 (P30- EY026877) to Stanford Ophthalmology (to all authors). The content is solely the responsibility of the authors and does not necessarily represent the official views of the National Institutes of Health.

