## Supplementary figure S1-S7 and Table S1-S2 for "Mettl14-mediated m^6^A modification ensures the cell cycle progression of late-born retinal progenitor cells" (from another version of this preprint: version 1, 2022-06-12)

### Supplementary information

1. Supplementary figures and figure legends (Figure S1-S7)

2. Supplementary tables (S1-S2)

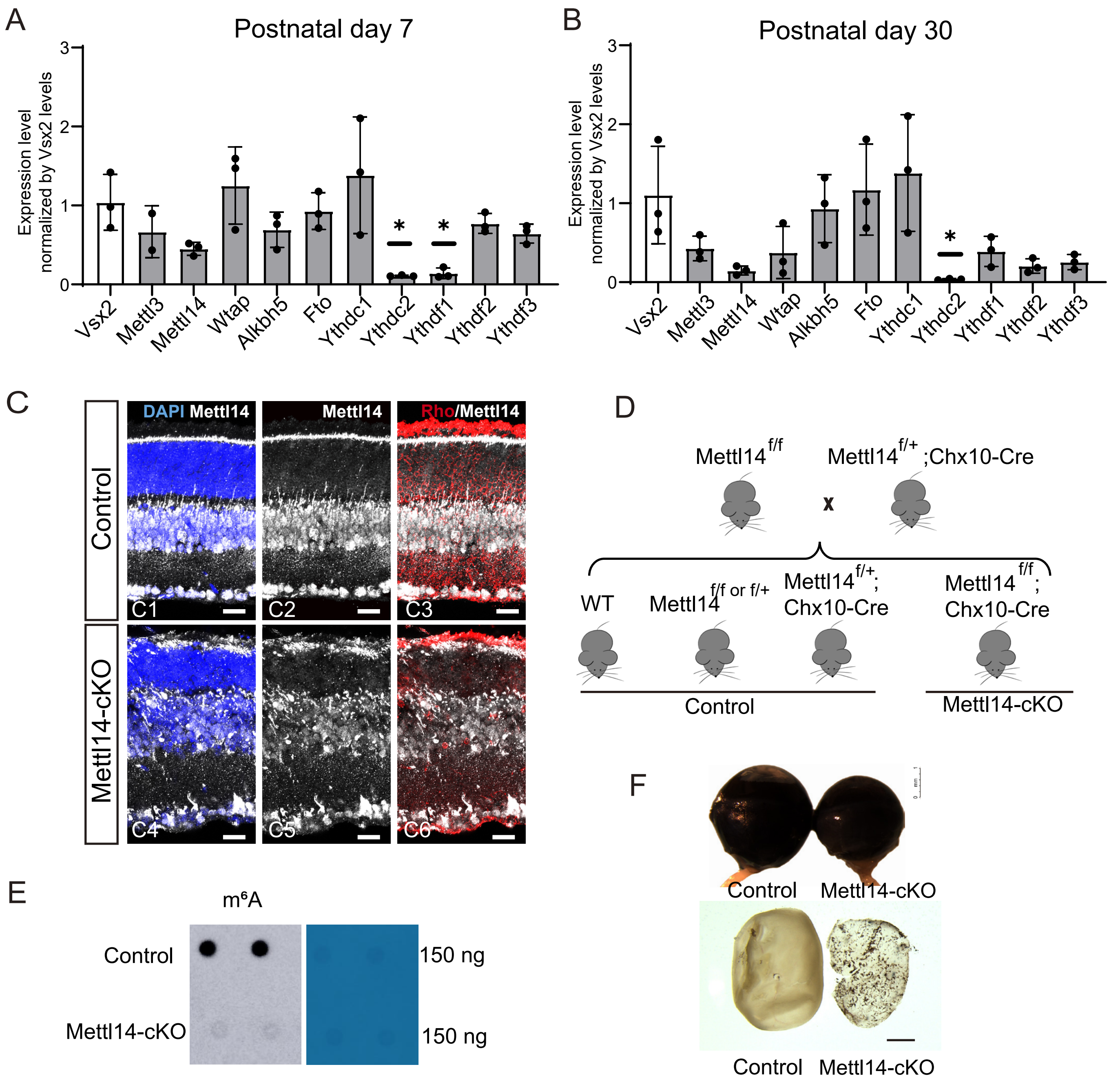
A
Postnatal day 7
Expression level normalized by Vsx2 levels
Vsx2
Mettl3
Mettl14
Wtap
Alkbh5
Fto
Ythdc1
Ythdc2
Ythdf1
Ythdf2
Ythdf3
B
Postnatal day 30
C
DAPI Mettl14
Mettl14
Rho/Mettl14
Control
Mettl14-cKO
C1
C2
C3
C4
C5
C6
D
Mettl14 f/f
Mettl14 f/+ ;Chx10-Cre
WT
Mettl14 f/f or f/+
Mettl14 f/+; Chx10-Cre
Mettl14 f/f; Chx10-Cre
Control
Mettl14-cKO
E
m6A
Control
Mettl14-cKO
150 ng
150 ng
F
Control
Mettl14-cKO
Control
Mettl14-cKO

**Figure S1. The expression of genes involved in m$^6$A modification in the retina and the gross morphology of Mettl14-cKO eyeballs in adults.**

**(A & B)** The expression levels of genes involved in m$^6$A modification in wildtype retinas at P7 (A) or P30 (B). All values were normalized to the levels of *Gapdh*. *Vsx2 (Chx10)*, which is abundantly expressed in the developing retina, serves as the control. Mean ± SD (n=3, * $P< 0.05$; Two tailed Student's T test). **(C)** The expression of METTL14 protein in control and Mettl14-cKO retinas at P21. Rho (Rhodopsin) marks the outer segments of rods. Scale bar: 20μm. **(D)** The breeding strategy for generating control and Mettl14-cKO mice. **(E)** The total levels of m$^6$A modification in control and Mettl14-cKO retinas at P7 measured by m$^6$A dot blot analyses. Methylene blue staining shows the loading amount of RNA (150ng). **(F)** The eyeballs (Top) and dissected retinas (Bottom) of adult control and Mettl14-cKO mice. Scale bar: 1mm.

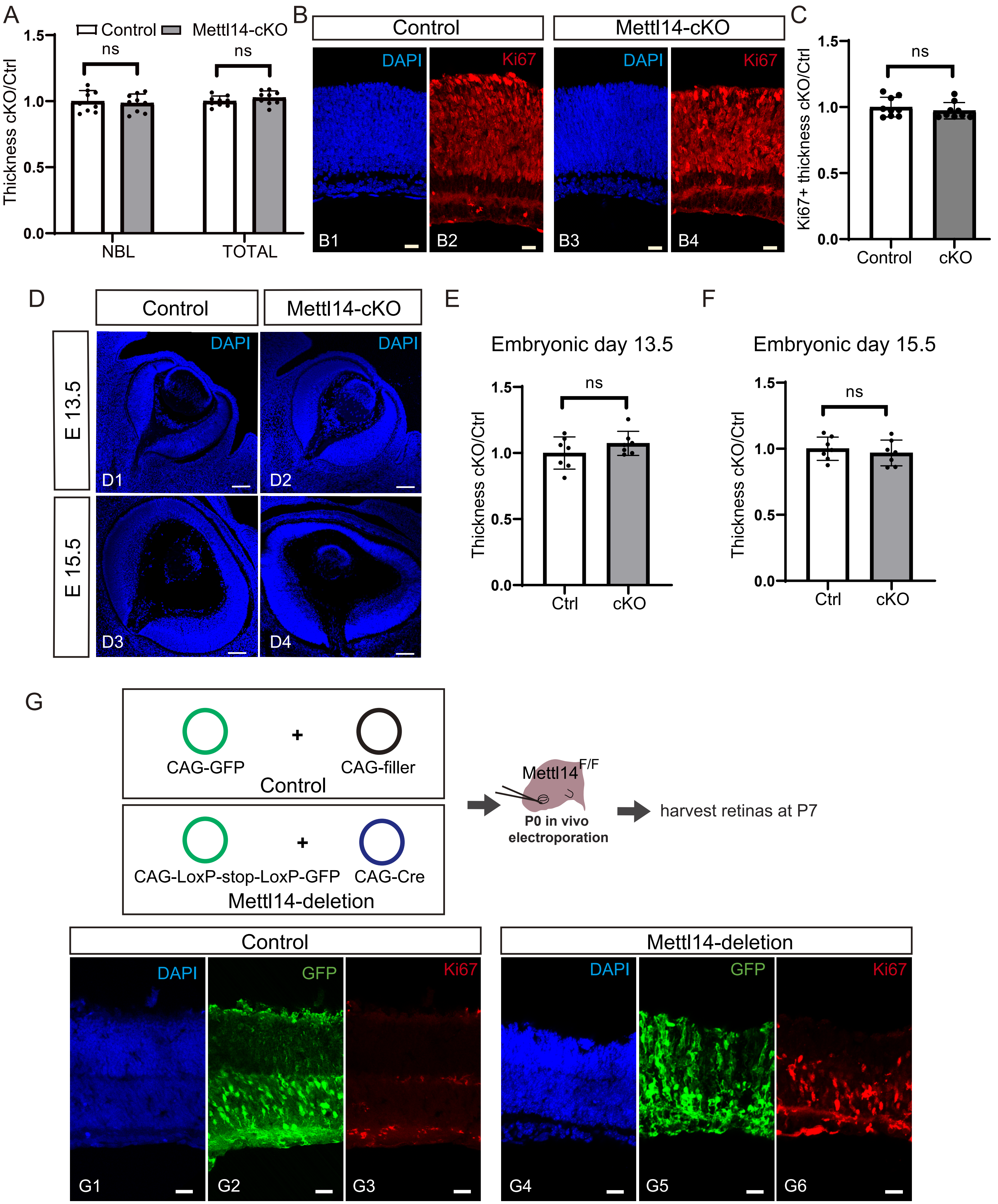

A
Control
Mettl14-cKO
ns
ns
Thickness cKO/Ctrl
1.5
1.0
0.5
0.0
NBL
TOTAL
B
Control
Mettl14-cKO
DAPI
Ki67
DAPI
Ki67
B1
B2
B3
B4
C
Ki67+ thickness cKO/Ctrl
ns
1.5
1.0
0.5
0.0
Control
cKO
D
Control
Mettl14-cKO
E 13.5
E 15.5
DAPI
DAPI
D1
D2
D3
D4
E
Embryonic day 13.5
Thickness cKO/Ctrl
ns
1.5
1.0
0.5
0.0
Ctrl
cKO
F
Embryonic day 15.5
Thickness cKO/Ctrl
ns
1.5
1.0
0.5
0.0
Ctrl
cKO
G
CAG-GFP
+
CAG-filler
Control
CAG-LoxP-stop-LoxP-GFP
+
CAG-Cre
Mettl14-deletion
Mettl14 F/F
P0 in vivo electroporation
harvest retinas at P7
Control
Mettl14-deletion
DAPI
GFP
Ki67
DAPI
GFP
Ki67
G1
G2
G3
G4
G5
G6

**Figure S2. The gross morphology of Mettl14-cKO retinas at or prior to birth and the effects of *Mettl14* deletion at P0.**

**(A)** The thickness of control and Mettl14-cKO retinas at P0. NBL: neuroblastic layer. Data were normalized to controls. Mean ± SD (n=9; Two tailed Student's T test). ns: not significant. **(B)** The Ki67 expression in control and Mettl14-cKO retinas at P0. Scale bar: 20μm. **(C)** The thickness of the Ki67+ regions in control and Mettl14-cKO retinas at P0. Data were normalized to control samples. Mean ± SD (n=9; Two tailed Student's T test). ns: not significant. **(D)** The control and Mettl14-cKO retinas at E13.5 and E15.5. Scale bar: 200μm. **(E & F)** The total thickness of control and Mettl14-cKO retinas at E13.5 (E) and E15.5 (F). Mean ± SD (n=9; Two tailed Student's T test). **(G)** Deletion of *Mettl14* by electroporating the CAG-Cre plasmids into *Mettl14*$^{F/F}$ mouse retinas at P0. The experimental design is shown on top. The retinas were harvest at P7 and stained with anti-Ki67 antibody (Bottom). Scale bar: 20μm.

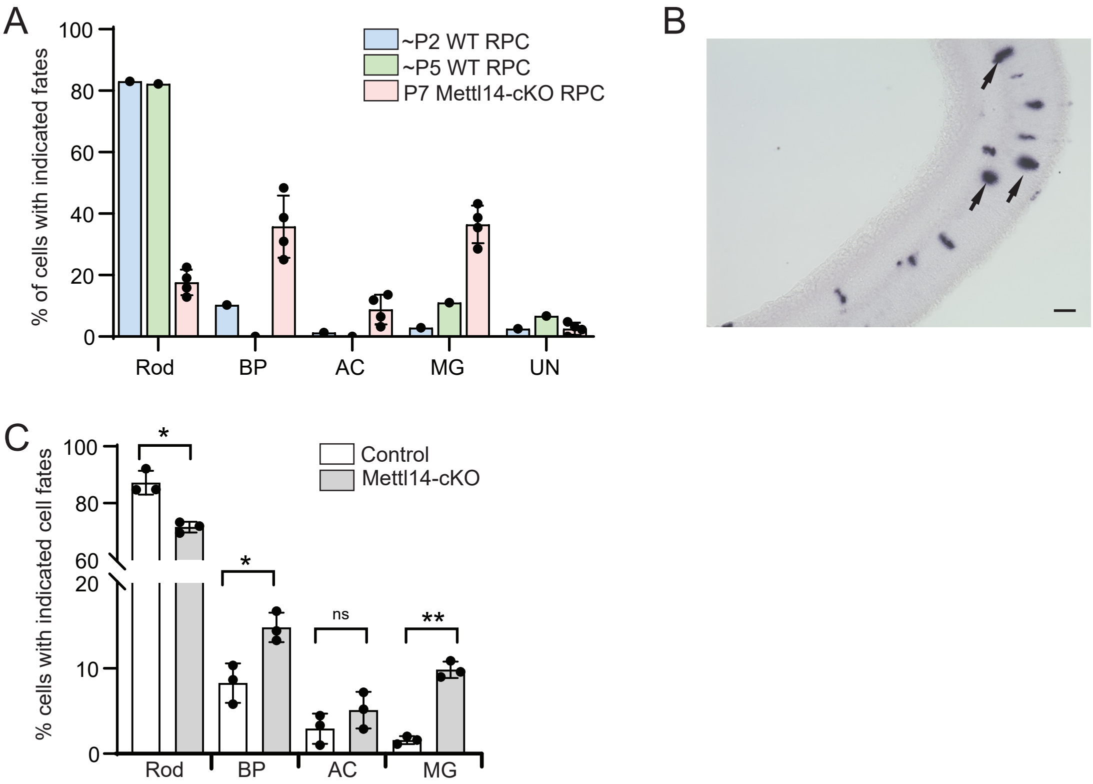
A
100
80
60
40
20
0
% of cells with indicated fates
~P2 WT RPC
~P5 WT RPC
P7 Mettl14-cKO RPC
Rod
BP
AC
MG
UN
B
C
*
*
ns
**
Control
Mettl14-cKO
100
80
60
20
10
0
% cells with indicated cell fates
Rod
BP
AC
MG

**Figure S3. The cell fates of P7 or P0 Mettl14-cKO RPCs.**

**(A)** The progeny cell fates of the P2, P5 WT RPCs and Mettl14-cKO P7 RPCs. The data of P2 and P5 RPCs were adopted from Turner et. al 1987. The data of Mettl14-cKO P7 RPCs were presented as mean ± SD (n=4). BP: bipolar cells; AC: amacrine cells; MG: Müller glial cells; UN: cells with unidentified fates. **(B)** The cells with unidentified fates (black arrows) in Mettl14-cKO retinas at P9. Scale bar: 200μm. **(C)** The progeny cell fates of P0 control and Mettl14-cKO RPCs. BP: bipolar cells; AC: amacrine cells; MG: Müller glial cells. Mean ± SD (n=3; *P< 0.05; **P<0.01; Two tailed Student's T test). ns: not significant.

A

| Before QC | Control | cKO |
|---|---|---|
| Total Cells | 13,287 | 15,319 |
| Total Reads | 199,704,911 | 223,690,818 |
| Total Genes | 21,407 | 21,473 |
| Mean Reads/Cell | 15,030 | 14,602 |
| Median Genes/Cell | 1,661 | 1,586 |
| Median UMI Counts/Cell | 3,062 | 2,887 |

| After QC | Control | cKO |
|---|---|---|
| Total Cells | 10,451 | 13,116 |
| Total Genes | 17,935 | 18,044 |
| Median Genes/Cell | 1,765 | 1,687 |
| Median UMI Counts/Cell | 3,072 | 2,885 |

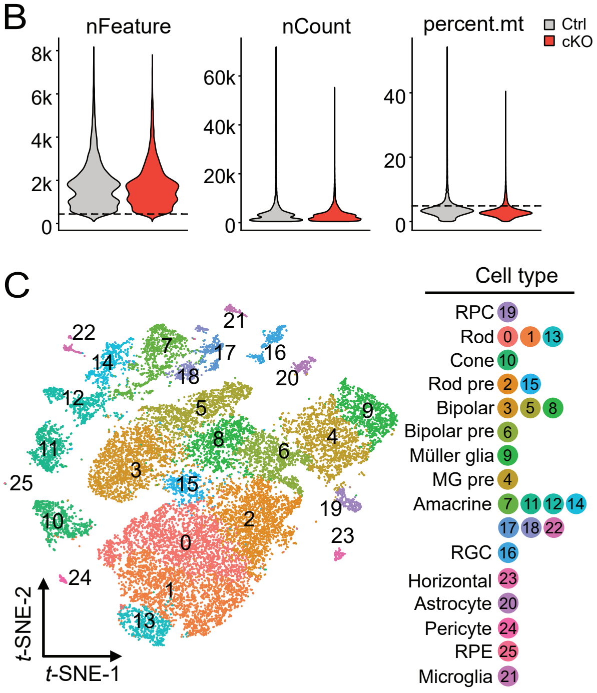


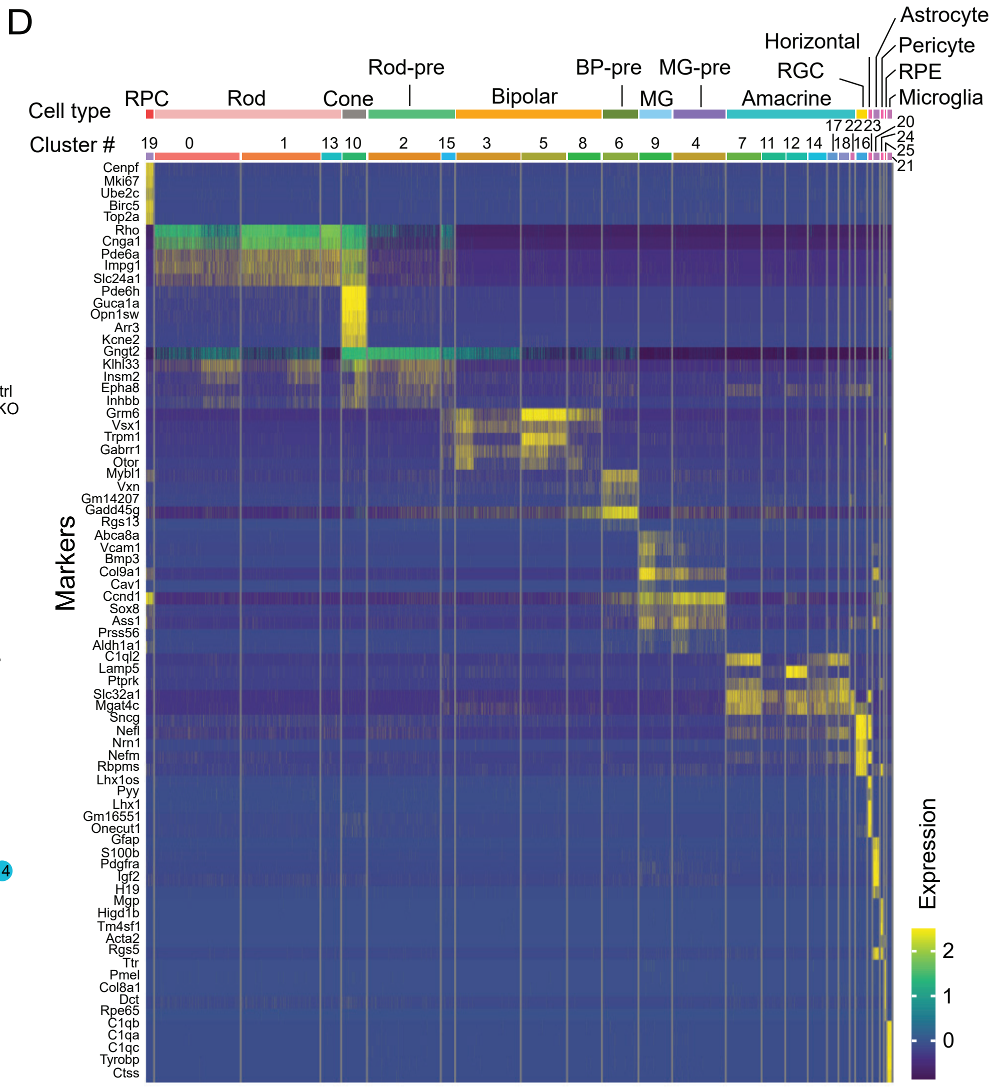

**Figure S4. Quality control and clustering of scRNA-seq data.**

**(A)** Summary of QC metrics for control and Mettl14-cKO samples. **(B)** Violin plots show the distribution of feature and count numbers, and percent of mitochondrial RNAs. The cutoff of QC is > 500 features/cell and < 5% reads as mitochondrial RNAs (shown as dash lines). **(C)** t-SNE plot representations of the 25 clusters (grouped by color) that were generated by unbiased clustering method (Seurat 4.1). These clusters were assigned to 15 cell types. **(D)** Heat plot shows the expression levels of the 5 top conserved genes for each cell type. The clusters are grouped by cell types.

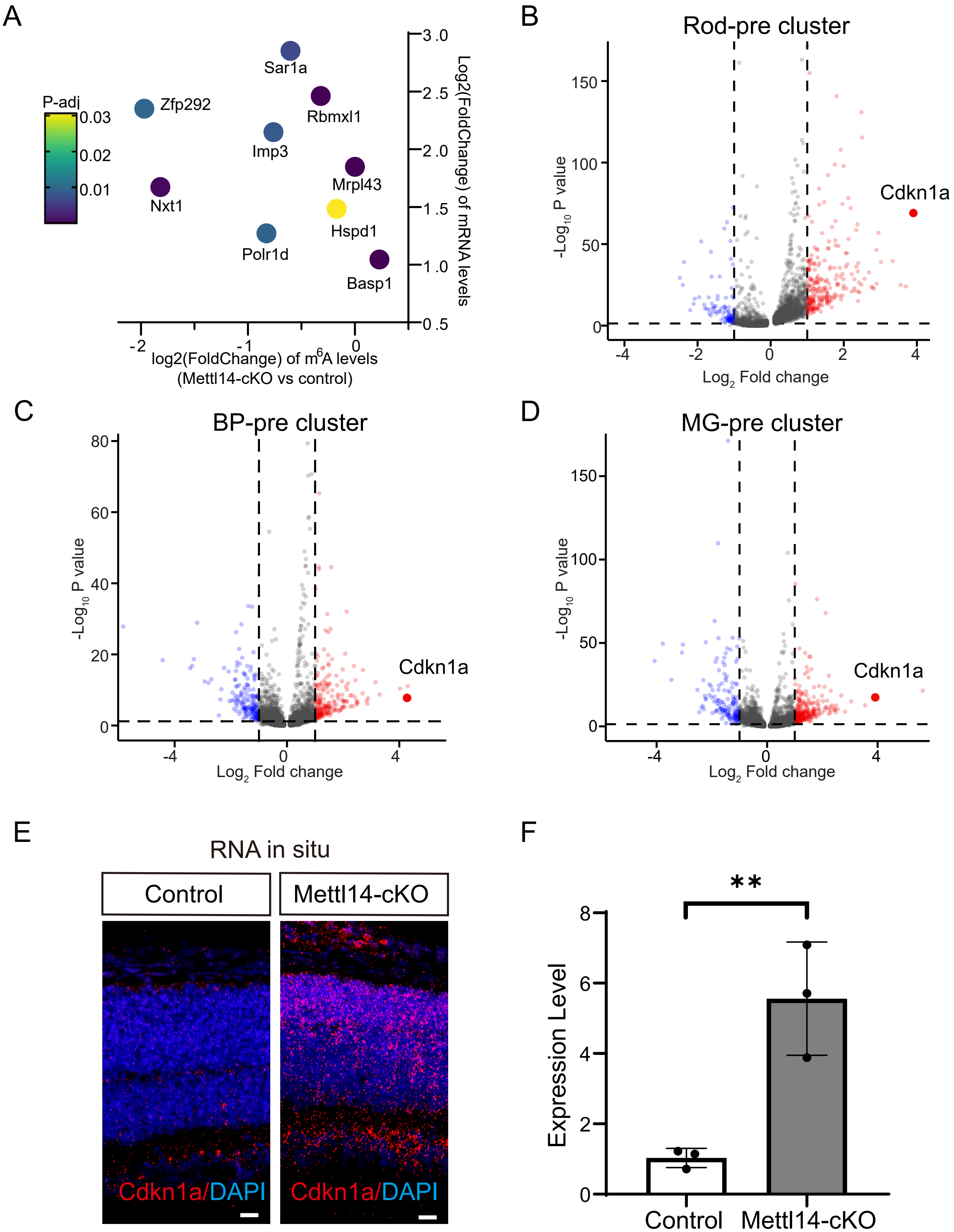
A
P-adj
0.03
0.02
0.01
Zfp292
Sar1a
Rbmxl1
Imp3
Mrpl43
Nxt1
Hspd1
Polr1d
Basp1
log2(FoldChange) of m6A levels
(Mettl14-cKO vs control)
Log2(FoldChange) of mRNA levels
B
Rod-pre cluster
Cdkn1a
-Log10 P value
Log2 Fold change
C
BP-pre cluster
Cdkn1a
D
MG-pre cluster
Cdkn1a
E
RNA in situ
Control
Mettl14-cKO
Cdkn1a/DAPI
F
**
Expression Level
Control
Mettl14-cKO

**Figure S5. scRNA-seq reveals the differentially expressed genes in Mettl14-cKO retinas at P7**

**(A)** The Bubble plot shows the $m^6A$ level changes and mRNA level changes of the top differential expressed genes (DEGs) in the RPC cluster (Mettl14-cKO vs control, $log_2$(FoldChange)>1, adjusted P value< 0.05 (Wilcoxon rank-sum test followed by Bonferroni correction for multiple comparisons). **(B-D)** The volcano plots show the DEGs in Rod-pre (B), BP-pre (C) and MG-pre (D) clusters (Mettl14-cKO vs control). The top candidate *Cdkn1a* (encodes P21 protein) is highlighted. Red: up regulated. Blue: down regulated. The cutoff for DEGs is P < 0.05 (Wilcoxon rank sum test) and **|**log2(FoldChange)**|** > 1 (dashed lines). **(E)** RNA *in situ* shows the expression of *Cdkn1a* in control and Mettl14-cKO retinas at P7. Scale bar: 20μm. **(F)** QPCR shows the levels of *Cdkn1a* mRNA in control and Mettl14-cKO retinas at P7. All values were normalized to the levels of *Gapdh*. Mean ± SD (n=3, ** P< 0.01; Two tailed Student's T test).

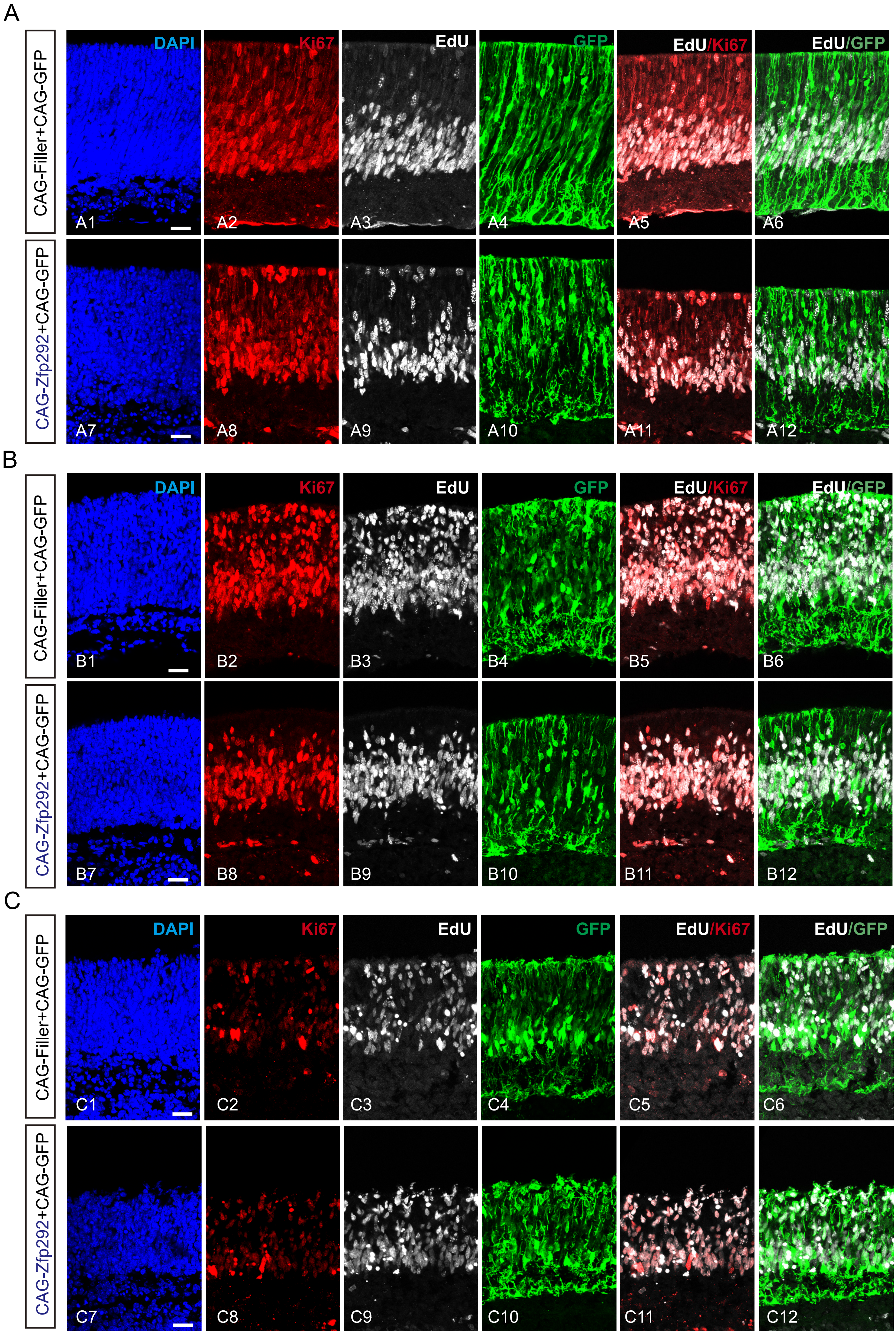
A
CAG-Filler+CAG-GFP
CAG-Zfp292+CAG-GFP
DAPI
Ki67
EdU
GFP
EdU/Ki67
EdU/GFP
A1
A2
A3
A4
A5
A6
A7
A8
A9
A10
A11
A12
B
B1
B2
B3
B4
B5
B6
B7
B8
B9
B10
B11
B12
C
C1
C2
C3
C4
C5
C6
C7
C8
C9
C10
C11
C12

**Figure S6. Over-expression of *Zfp292* prevents RPCs from exiting the cell cycle.**
**(A-C)** Ki67 staining and EdU detection of control (*CAG-filler*+*CAG-GFP*) and *Zfp292*-overexpressing (*CAG-Zfp292*+*CAG-GFP*) retinas at 2 hours (A), 30 hours (B) or 60 hours (C) after EdU pulse (1 hour pulse).

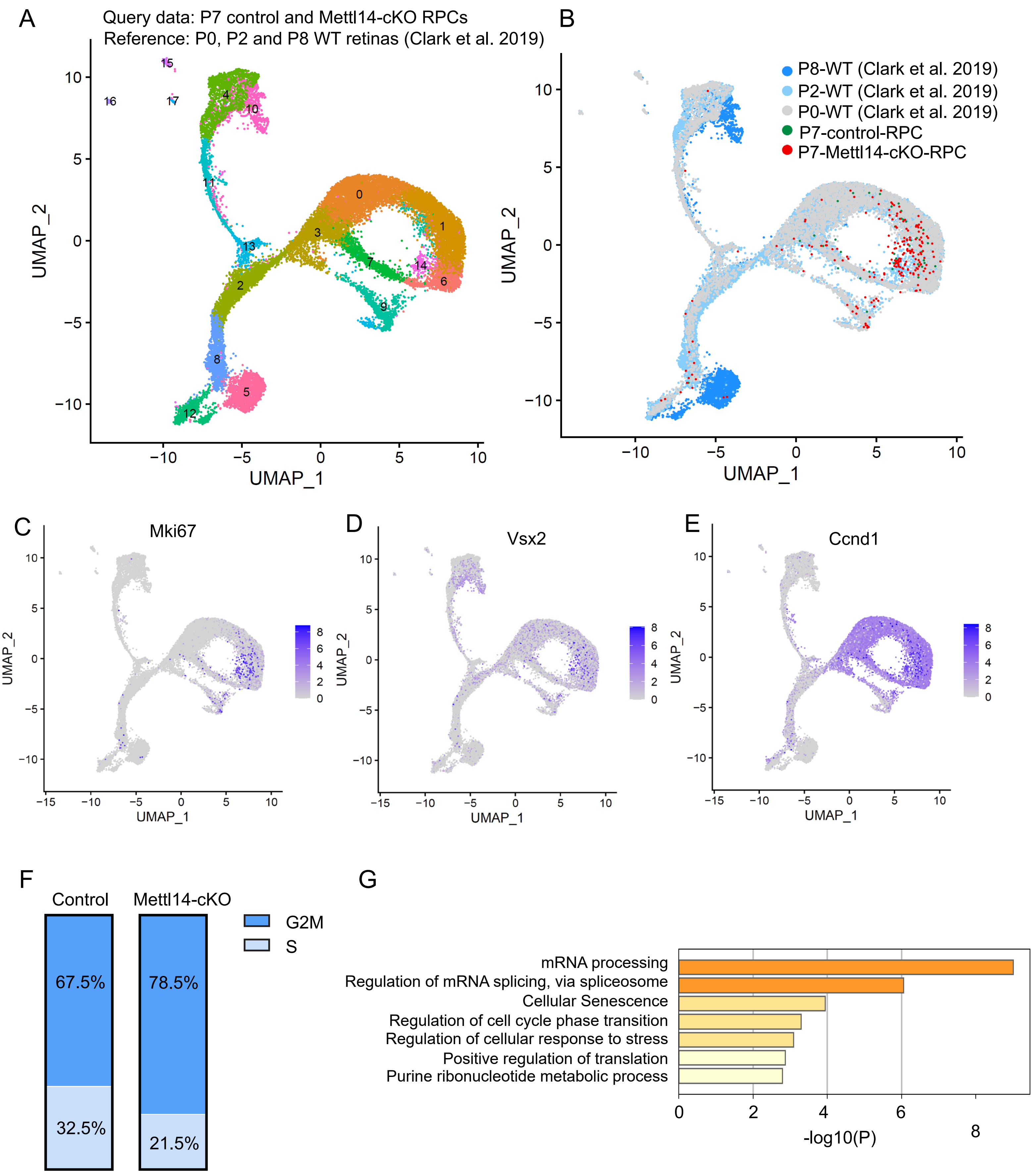
A
Query data: P7 control and Mettl14-cKO RPCs
Reference: P0, P2 and P8 WT retinas (Clark et al. 2019)
UMAP_1
UMAP_2
B
P8-WT (Clark et al. 2019)
P2-WT (Clark et al. 2019)
P0-WT (Clark et al. 2019)
P7-control-RPC
P7-Mettl14-cKO-RPC
UMAP_1
UMAP_2
C
Mki67
D
Vsx2
E
Ccnd1
F
Control
Mettl14-cKO
G2M
S
67.5%
32.5%
78.5%
21.5%
G
mRNA processing
Regulation of mRNA splicing, via spliceosome
Cellular Senescence
Regulation of cell cycle phase transition
Regulation of cellular response to stress
Positive regulation of translation
Purine ribonucleotide metabolic process
-log10(P)

**Figure S7. Comparison of P7 Mettl14-cKO PRCs with P0, P2, and P8 wildtype RPCs. (A)** UMAP plot shows the 17 clusters (grouped by color) that were generated by querying the P7 Mettl14-cKO RPCs in the reference dataset (combined P0, P2, and P8 retinal data from Clark et al. 2019). **(B)** UMAP plot shows the cells from different samples. **(C-E)** UMAP plots show *Mki67*+ (C), *Vsx2*+ (D), and *Ccnd1*+ (E) RPCs cells in the query and reference combined dataset. **(F)** Seurat cell cycle analyses of control and Mettl14-cKO RPCs at P7. G2M: G2 and M phases of the cell cycle; S: S phase of the cell cycle. **(G)** Gene Ontology analysis (metascape.com) of the genes with increased mRNA levels in P7 Mettl14-cKO RPCs compared to P0, P2 and P8 WT RPCs (adjusted P value<0.05, log2 (FC)>1.5).

**Table S1 Primers used in this study**

| Name | Sequence (5'-3') | Experiments |
|---|---|---|
| GAPDH-f | ACCACAGTCCATGCCATCAC | qPCR |
| GAPDH-r | TGGACCACCCTGTTGCTGTA | qPCR |
| Mettl14-f | TCCTCCTGCTGCATTTCCAGTTC | qPCR |
| Mettl14-r | TGCTGAAACCAGAGAAACCTGCA | qPCR |
| Vsx2-f | CCAGCTCCGATTCCGAAGATGTT | qPCR |
| Vsx2-r | TGAATGCTTTCTCCAGCTCCTCT | qPCR |
| Mettl3-f | TGGAGCTCTATCCAGGCCCATAA | qPCR |
| Mettl3-r | CGTAGAGATGGCAAGACGGATGG | qPCR |
| WTAP-f | GGCTGCTGAACTTGCTTGAGGTA | qPCR |
| WTAP-r | CGATGTGACTGGTTTAAGGGAATCTG | qPCR |
| ALKBH5-f | CCCATCGTGTCCGTGTCTTTCTT | qPCR |
| ALKBH5-r | TCATCAGCAGCATACCCACTGAG | qPCR |
| FTO-f | CACTTGGCTTCCTTACCTGACCC | qPCR |
| FTO-r | TATGCAGTGTGAGAAAGGCCTCG | qPCR |
| YTHDC1-f | TGTCTCGTGGTCAGAGCCATAGT | qPCR |
| YTHDC1-r | TCTGAAAGAAGGGCCAAGTCTC | qPCR |
| YTHDC2-f | CCTGTGAAAAGGGGAGAGTCTGAA | qPCR |
| YTHDC2-r | AGGAACTGAGGAATCTGTGTGGT | qPCR |
| YTHDF1-f | TCCGAGTAACAGTTACCCCTCGA | qPCR |
| YTHDF1-r | CATGCATGAAGTGATGGTCTCCA | qPCR |
| YTHDF2-f | GTAGAAAGTGGGGCTCTCCGTTG | qPCR |
| YTHDF2-r | GCCTCTTGGAGCAGAGACCAAAA | qPCR |
| YTHDF3-f | CGTGGATCAGAGACCTAAAGGGC | qPCR |
| YTHDF3-r | TGGAAATCCAATGGACGGAGCAT | qPCR |
| Cdkn1a-f | CCCGTGGACAGTGAGCAGTTG | qPCR |
| Cdkn1a-r | GGTAGACCTTGGGCAGCCCTAG | qPCR |
| Dbf4-r | CCATGAGGATCCACAGCAAAGCA | qPCR |
| Dbf4-f | TACAGGAGAAACGCGACCCAATG | qPCR |
| Tpx2-r | GCGAGCCTTTTCAGGGGAAGAAT | qPCR |
| Tpx2-f | CTCTCTTAGCATCCAGGGAAGAACA | qPCR |
| Foxm1-f | AAAGATGAGTTCTGACGGGCTGG | qPCR |
| Foxm1-r | AAGAGTAGGGTGGCCTCTCAGAC | qPCR |
| Ntrk3-f | GGAACCTCTTCCCTCTCCTGGAA | qPCR |
| Ntrk3-r | GAGCTCCATGTCCACAGCATTGA | qPCR |
| Birc5-f | TTCATCCACTGCCCTACCGAGAA | qPCR |
| Birc5-r | TGCTTCTTGACAGTGAGGAAGGC | qPCR |
| Ttk-f | AGTTAATTGGCAGCAGTGTGACG | qPCR |
| Ttk-r | GCAAAGCTCTCATTCTGGCCGTA | qPCR |
| fbxo5-f | CACGCTGACTGGAGGACTTGATT | qPCR |
| fbxo5-r | GCAAAACCATGCAGCGAGTCATT | qPCR |
| Brip1-f | TGGCAGATAACTTTGCAGCCAGA | qPCR |
| Brip1-r | CTACAGCGACACAAACCTGGACA | qPCR |

| | | |
|---|---|---|
| Brca2-f | ACCTTCCCTAATTCTCTGAAAGGCG | qPCR |
| Brca2-r | AGTATAAGCCCCACGGTTATGAACC | qPCR |
| Zfp292 amp- primer1 | GTCTCATCATTTTGGCAAAGATGGCtGAtGATGaaGCtGA | Cloning |
| Zfp292 amp- primer2 | ATGGCtGAtGATGaaGCtGA | Cloning |
| Zfp292 amp- primer3 | TACGTATTATAATGTTGGGC | Cloning |
| Zfp292 amp- primer4 | GCCCAACATTATAATACGTA | Cloning |
| Zfp292 amp- primer5 | CCCACTTATAACAGAAACAT | Cloning |
| Zfp292 amp- primer6 | CAAAATTTTACCAATAATAG | Cloning |
| Zfp292 amp- primer7 | GGTTGTGAtTCAGTcTTTGT | Cloning |
| Zfp292 amp- primer8 | ACAAAgACTGAaTCACAACC | Cloning |
| Zfp292 amp- primer9 | atttacgtagcggccgcGATTCAGTACTGCCCTCTGCCCG | Cloning |
| Zfp292 amp- primer10 | TCAGTACTGCCCTCTGCCCG | Cloning |
| YNST-cdkn1a-A1P1E | AGACGTTTAAATTAAAACACAAATAaaCGTCAACGACAAGC | RNA in situ |
| YNST-cdkn1a-A1P1O | AGAGAATCATACGTtaAATTAAGACACACAGAGTGAGGGCT | RNA in situ |
| YNST-cdkn1a-A1P2E | GCCGAAGATGGGGAAGAGGCCTCCTaaCGTCAACGACAAGC | RNA in situ |
| YNST-cdkn1a-A1P2O | AGAGAATCATACGTtaCCCACAGCAGAAGAGGGCGGGGCTC | RNA in situ |
| YNST-cdkn1a-A1P3E | CCCCACACTAAGGGCCCTACCGTCCaaCGTCAACGACAAGC | RNA in situ |
| YNST-cdkn1a-A1P3O | AGAGAATCATACGTtaCTAATTTAGGTTTGTTTTGTTTTGG | RNA in situ |
| YNST-cdkn1a-A1P4E | TTGAGTCCTAACTGCCATCCCTGTTaaCGTCAACGACAAGC | RNA in situ |
| YNST-cdkn1a-A1P4O | AGAGAATCATACGTtaAGGCTGTGACTGCTTCACTGTCATC | RNA in situ |
| YNST-cdkn1a-A1P5E | TATAGGAAACACAGAGCTTGGGTTGaaCGTCAACGACAAGC | RNA in situ |
| YNST-cdkn1a-A1P5O | AGAGAATCATACGTtaAGGGGCTTAAATAATAATAATCTAC | RNA in situ |
| YNST-cdkn1a-A1P6E | CAGAGACCCACAGGAGAGGTGGGCTaaCGTCAACGACAAGC | RNA in situ |
| YNST-cdkn1a-A1P6O | AGAGAATCATACGTtaCCAGGAGTGGCAAAGGGGATCCAGA | RNA in situ |
| YNST-cdkn1a-A1P7E | CTGCATATACATTCCCTTCCAGTCCaaCGTCAACGACAAGC | RNA in situ |
| YNST-cdkn1a-A1P7O | AGAGAATCATACGTtaTGAGCTGTGGGGCAAGTGCCTAGAT | RNA in situ |
| YNST-cdkn1a-A1P8E | CCCCACCACCACACACCATAGAATGaaCGTCAACGACAAGC | RNA in situ |
| YNST-cdkn1a-A1P8O | AGAGAATCATACGTtaCTGGGAGGCCTGTCTCACCACCAAG | RNA in situ |
| YNST-cdkn1a-A1P9E | AATCATCGAGAAGTATTTATTGAGCaaCGTCAACGACAAGC | RNA in situ |
| YNST-cdkn1a-A1P9O | AGAGAATCATACGTtaCAGCTTTGGGGTCGGGTGTGAGGAC | RNA in situ |
| YNST-cdkn1a-A1P10E | AGGGAGGGAGCCACAATACATGTTCaaCGTCAACGACAAGC | RNA in situ |
| YNST-cdkn1a-A1P10O | AGAGAATCATACGTtaGGGGTGCGGGCTAAGGGTAGACAGT | RNA in situ |

**Table S2 The sequences of gblocks used for constructing *CAG-Zfp292***

| | Sequencing (5'-3') |
|---|---|
| Zfp292 g-block1 | ATGGCtGAtGATGaaGCtGAaCAaGAaCGaCTtAGtGGTGGaGGa<br>TGtGCtGCtGAaCTtCGaCGaCTtGGaGAaCGtCTGCAAGAGCTG<br>GAGCGaCGaCTATGCGAGAGCCGtGAGCCaGCtGTGGAGGCa<br>GCaGCtGCCTACTGCCGCCAGCTGTGCCAGACACTTCTTGAA<br>TATGCAGAGAAATGGAAgACcTCCGAAGACCCTTTACCTTTGC<br>TGGAGGTATACACAGTGGCaATCCAGAGCTACGTTAAAGCTC<br>GACCATACCTTACCTCTGAGTGTGAAAGTGTAGCCTTGGTGC<br>TGGAACGCTTGGCATTAAGtTGcGTTGAACTTTTACTGTGTCT<br>GCCTGTTGAGCTATCAGATAAGCAGTGGGAACAATTcCAGAC<br>CTTGGTACAGGTAGCGCATGAGACACTGATGGAGAGTGGCA<br>GCTGTGAACTGCAGTTCCTAGCTACTCTAGCTCAGGAGACTG<br>GGGTGTGGAAAAACGCAGTGCTCTCCACTATCCTTTCCCAGG<br>AACCACTGGATAAAGAGAAAGTGAATGAATTTTTAGCTTTTGA<br>GGGTCCCATCTTGTTGGATATGAGAATTAAACATCTAATCAAA<br>ACCAATCAGTTAAGTCAAGCAACCGCTCTAGCAAAaCTGTGcT<br>CTGATCATCCTGAGATTGGTACAAAAGGTAGTTTTAAACAAAC<br>TTACCTTGTCTGTCTTTGtACATCATCTCCAAGTGAgAAGCTAA<br>TTGAAGAGATcTCAGAAGTGGATTGCAAAGATGCGTTAGAAAT<br>GATCTGTAACTTAGAGTCAGAGGGTGATGAGAAAAGCGCCCT<br>GGTcTTGTGTACTGCtTTTCTcTCACGGCAGCTCCAGCAAGGA<br>GATATGTACTGTGCTTGGGAACTCACTCTGTTTTGGAGTAAAT<br>TGCAGCAAAGGGTGGAACCATCAGTACAAGTGTACCTGGAAA<br>GGTGTCGaCAGCTTTCCCTaTTAACAAAGACTGTGTATCACAT<br>TTTCTTtCTGATTAAAGTTATTAATTCAGAGACTGAAGGGGCTG<br>GACTCGCTACCTGCATAGAATTATGTGTAAAAGCTCTTCGCTT<br>GGAATCTACAGAAAATACTGAAGTGAAAATATCAATTTGCAAA<br>ACCATTTCATGTTTGTTGCCTGAGGATCTGGAAGTCAAACGT<br>GCTTGTCAGTTGAGTGAATTTCTTATTGAGCCaACAGTAGATG<br>CATATTATGCTGTGGAGATGTTATACAACCAGCCAGACCAGA<br>AATATGATGAAGAGAATCTTCCAATACCAAATTCTCTACGTTG<br>TGAGCTCTTACTTGTTTTGAAAACTCAGTGGCCCTTTGATCCA<br>GAgTTTTGGGATTGGAAAACTTTGAAGCGCCAGTGTCTTGCT<br>CTCATGGGAGAAGAGGCGTCTATTGTGTCCTCAATTGATGAA<br>CTGAATGACAGTGAAGTCTATGAGAAAGTAGACTACCAGGGT<br>GAAAGGGGAGACACATCTGTGAATGGaCTTTCaGCTGCTGGA<br>CTTGGTACTGATTCTGGCCTGCTGATGGATACTGGTGAcGAA<br>AAGCAGAAGAAGAAAGAGATAAAAGAgTTgAAAGATAGGGGG<br>TTTATATCTGCTAGGTTTAGGAATTGGCAAGCCTACATGCAGT<br>ATTGTTTGCTCTGTGACAAgGAATTCCTTGGACACAGAATAGT<br>ACGGCATGCTCAAAAACATTACAAAGATGGGATTTACAGCTG<br>TCCCATATGTGCAAAGAATTTcAATTCTAAAGACTCGTTTGTC<br>CCTCATGTTACCCTGCATGTTAAACAGTCTAGTAAAGAGAGA<br>CTAGCAGCTATGAAGCCATTAAGAAGATTGGGAAGGCCTCCT<br>AAAATCACAGCCACCCATGAAAATCAgAAGACTAATATTAATA |

| | |
|---|---|
| | CaGTGGCTAAACAGGAACAGCGACCCATAAAAAAGAATAGcC<br>TTTATTCAACAGATTTCATAGTGTTTAATGACAACGATGGTTC<br>AGATGATGAAAAcGACGACAAGGACAAGTCgTATGAGCCCGA<br>GGTGATCCCTGTCCAGAAACCAGTACCTGTTAATGAGTTTAAT<br>TGTCCTGTGACCTTTTGTAAAAAGGGCTTTAAGTACTTCAAAA<br>ATTTAATTGCTCATGTGAAAGGCCATAAGGATAGTGAAGATG<br>CCAAACGCTTTCTTGAAATGCAAAGCAAGAAAGTCATTTGCCA<br>GTACTGTAGAAGGCACTTTGTAAGCGTCACTCATCTCAATGAT<br>CACTTACAAATGCACTGTGGCAGTAAGCCATATATATGTATAC<br>AGATGAAATGTAAGGCTGGTTTTAATAGTTACGCAGAGCTGTT<br>AGCCCACCGAAAGGAGCAcCAAGTCTTTAGAGCAAAGTGCTT<br>ATTTCCAAAATGTGGCAGAATTTTTTCCCAAGCTTATTTACTGT<br>ATGATCATGAGGCCCAACATTATAATACGTA |
| Zfp292 g-block2 | GCCCAACATTATAATACGTACACGTGCAAGTTCACAGGTTGT<br>GGTAAGGTGTACCGTTCTCAGAGCGAGATGGAGAAGCACCA<br>GGATGGCCACAGTCATCCTGAAACAGGGCTGCCTCCTGAAG<br>ACCAGCTTCAGCCATCTGGAAATGATGTGAATCCGGACTCAG<br>GAGCGACGGCTGCAGGAGGAAGGTCCGAGAACAGCATTGAC<br>AAGAACCTGGGTTCAAACAGAAGTGCtGATTGGGAGAAAAAC<br>AGAGCAGAGCCAGCTGTGACTAAACACGGCCAGATCTCTGC<br>CGCTGAACTCAGGCAAGCTAACATACCATTGTCAAATGGTCT<br>GGAAACCCGTGATAATACTACTGTTCTTCGGACCAATGAAGT<br>AGCTGTGTCCATCAAGGTGTCTGTCAAtCATGGGGTAGAGGG<br>TGACTTTGGAAAGCAAGAAAACCTAACCATGGAAGGCACTGG<br>TGAGcCGCTGATCACAGATGTGCATAAACCAGGTATAGGTGC<br>TGGGGTCCAGTTATGTCATCCAGGTTTtCAAGAgAAGAAgGGT<br>CACGAGTGCCTGAACGAAGCCCAGAATTCTTTATCAAACTCA<br>GAATCACTGAAGATGGATGACCTaAACCCACAAAGCTTAGAA<br>AGACAGGTGAACACTCTGATGACCTTcTCTGTACAAAATGAG<br>GCAGGACTTGAAGACAATTCtCAAATTTGCAAGTTTGAATGTG<br>GAGGTGATGTTAAAACCTCATCCAGCCTTTATGAcTTACCTCT<br>TAAGACACTAGAAAGTATCACATTTGTTCAGTCACAGCCCGA<br>CCTAAGCAGTCCGTTGGGATCTCCATCAGTACCTCCAAAAGC<br>TCCAGGaCAGAAGTTCAGCTGCCAGGTTGAGGGATGCACTC<br>GAACATATAACTCTTCACAGAGTATTGGAAAACACATGAAGAC<br>AGCACACCCTGACCAATATGCTGCTTTTAAACTGCAGCGCAA<br>GACGAAAAAAGGTCAGAAATCTAACAACTTAAATACACCAAAT<br>CAcGGAAAaTGTGTTTATTTTTTGCCATCACAAGTGAGCAGCT<br>CTAATCATGCTTTTTTTACACCACAGACCAAAGCCAAcGGGAA<br>CCCTGCCTGTTCAGCCCAGGTGCAGCATGTCTCGCCTTCCAT<br>TTTCCCAGCTCATTTAGCAAGTGTATCAGCTCCATTGTTACCC<br>TCAGTGGAAAGTGTCCTAAGTCCAAATATACCTTCTCAGGATA<br>AACATGGACAAGATGGCATATTATGTTCACAgATGGAAAATTT<br>GTCTAATGCTCCCTTGCCAGCACAAATGGAAGATCTAACCAA<br>AACAGTaTTGCCTTTGAATATTGACAGCGGCTCAGATCCGTTT<br>CTcCCTTTACCCACAGAAAATAGCTCTCTCTTCTCTTCACCAG |

| | |
|---|---|
| | CAGACAGTGAGAATAATTCTGTTTTTTCCCAACTGGAAAATAG TACAAATCATTATCCCTCCCAGACGGATGGAAACATAAATTCC TCTTTTCTGAAAGGAGGCAGCAGTGAgAATGGAGTTTTTCCTT CaCAAGTAAGTTCTGCAGATGACTTCAGTAGCACtAGTGCCCA ACCGTCTACACCTAAGAAAGTGAAAAAAGACCGTGGTCGAGG CCCAAATGGGAAGGAAAGgAAACCCAAGCACAACAAgAGGGC TAAATGGCCTGCGATTATCAGAGATGGGAAATTCATCTGTAG CAGGTGTTACAGaGCTTTCACCAACCCCAGGTCCCTGGGTGG ACACCTGTCTAAAAGGTCTTACTGCAAACCACTGGATGGAGC AGAAATAGCACAGGAACTTCTACAGACtAACAGACAGCCTTCC CTCCTAGCTAGCATGATTCTCTCCACAAGTGCAGTAAATATGC AACAGCCGCAACAGTCTAAtTTTAATCCAGAAACATGCTTTAA AGACCCATCATTCCTGCAACTTCTTAAcGTGGAAAATCGTCCA ACCTTTTTACCtAGTACATTTCCAAGATGTGACGTGAGTAACTT TAATGCCAGTGTTAGTCAGGAAGGaAGTGAAATTATTAAGCA GGCTTTAGAAACTGCTGGCATTCCCAGCACGTTTGAGAGTGC CGAAATGCTTTCTCAGGTTGTTCCAATAGGCAGTGTCTCCGA TGCAGCACAAGTCAGTGCAGCGGGGATGCCAGGGCCACCTG TGACACCCTTGTTACAGACTGTTTGCCACCCAAACACCTCAC CATCAAACCAGAATCAAACGCCAAATTCCAAAACCCTCAAAG AATGTAACAGTTTGCCTCTCTTTACAACAAATGATTTACTGCT AAAGACTATTGAAAAcGGCTTGTGCTCCAATTCATTCAGTAGT TCTACTGAACCACCACAAAATTTTACCAATAATAGTGCACATG TTTCTGTTATAAGTGGG |
| Zfp292 g-block3 | CAAAATTTTACCAATAATAGTGCACATGTTTCTGTTATAAGTG GGCCTCAGAATACAAGATCCAGTCATTTGAATAAAAAAGGAA ATAGTGCATCTAAGAAGAGAAAgAAAGTTGCTCCTGCAGTAA GTGTATCTAATACTTCCCAAAATGTGCTACCAACTGATTTACC AGTGGGCCTTCCAGCAAAGAATCTTCCAGTCCCTGATACCAA CACACGGTCAGACATGACTCCAGATTGTGAACCTCGGGCTTT GGTGGAAAATCTCACACAGAAATTAAATAACATTGACAATCAT TTGTTTATAACTGATGTAAAAGAGAACTGTAAAGCCAGTCTTG AGCCCCATACAATGTTAACCCCTTTAACATTAAAAACGGAAAA CGGCGATTCCCGAATGATGCCTTTGAGTTCATGCACACCAGT GAATTCTGATTTGCAGATTTCTGAAGATAATGTTATTCAGAAC TTTGAGAAGACTCTTGAAATTATTAAAACTGCTATGAATTCTCA AATACTTGAGGTAAAAAGTGGATCTCAGGGTACTGGTGAGAC AACACAGAATGCTCAGATAAATTACAGCATGCAACTTCCCTCA GTAAACTCTATCCCAGATAACAAGCTGCCTGATGCTTCTCAGT GCTCCTCTTTCCTAACTGTAATGCCAACAAAGTCTGAAGCATT ACATAAGGAGGATCAAATACAGGACATTTTAGAGGGTTTGCA AAACTTAAAACTAGAAAATGACACCTCTGCTCCAGCTTCCCAG AGTATGCTAATGAACAAATCAGTAGCACTGTCCCCTACTCCTA CTAAATCAACTCCAAATATTGTAGTCCAGCCAGTACCCGAAGT GATACATGTTCAGCTTAATGACAGAGTTAATAAGCCCTTTGTG TGTCAAAACCAAGGCTGTAACTACAGTGCTATGACAAAGGAT |

| | |
|---|---|
| | GCCCTGTTTAAACACTATGGTAAAATCCATCAGTATACTCCAG AGATGATTCTTGAAATTAAGAAGAATCAATTAAAATTcGCTCCA TTTAAATGTGTAGTACCTTCATGTACCAAAACATTTACAAGAA ATTCTAATCTCCGGGCACACTGTCAGTTGGTGCATCATTTTAC AATAGAAGAAATGGTAAAGCTAAAAATAAAAAGGCCCTATGG AAGAAAATCTCAGAGTGAAAATTTGTCATCTCCACAGAATAAT CAAGTGAAGAAACAGCCATCCATGGCCGAGGAAACAAAgACT GAaTCACAACC |
| Zfp292 g-block4 | ACAAAgACTGAaTCACAACCAGCCTTtAAGGTACCAGCAGCAA CAGGTGATGCTGCACTTGCTAAcGCAACAGTAATCCCAGAAA AACAACTTGCAGAgAAAAAAAGTCCTGAGAAACCAGAAAGTTC TTCACAGCCTGTCACATCaTCTGCTGAACAATATAATGCAAAT CTcGCAAACCTAAAgACCAAgGGAAGGAAAAAcAAGAGGCATA GgAAgGAgAAGGAAGAgAAgCGGGAgAAGAAcCCAGTTTCCCA GGCCTTTGAACTTCCAACAAAgTACAGTTCGTACAGACCTTAC TGCTGTGTCCACCAGGGATGCTTTGCTGCaTTcACAATtCAGC AAAAtTTGATTCTTCATTAtCAGGCTGTACATAAgTCAAATCTTC CTACATTTTCTGCtGAGGTTGAAGAaGAgAGTGAgGCTGTTAAA GAAAGTGAAGAAACTGAACCGAAACAATCAATGAAAGAATTTA GGTGTCAGGTGAGTGACTGcTCTAGGATTTTCCAAGCAATTA CTGGCCTAATACAGCACTACATGAAACTTCATGAAATGACCC CCGAGGAAATTGAgAGCATGACTGCaGCTGTaGATGTTGGCA AATTcCCATGTGATCAGTTGGAGTGTAAGTTaTCTTTTACAACA TACCTGAGCTATGTTGTTCATCTTGAGGTAGACCATGGAATTG GAACAAGGACAAGcAAGGCtGAAGAAGAcGGCATATACAAaTG TGACTGTGAGGGCTGTGACAGGATATATGCCACTCGGTCTAA TCTTCTCCGACACATCTTTAATAAgCATAATGACAAGCATAAA GCCCATCTGATTCGGCCAAGAAAgTTAACTGGCCAGGAAAAc ATATCAAGTAAGGCAAACCAgGAgAAATCAAAGTCTAAgCATC GGACAACAAAACCCAACAGATCCGGGAAAGACGGAATGAAA ATGCCAAAGACtAAGCGAAAGAAgAAgAGTAATTTAGAAAACA AGAGCGCAAAAGTAGTGCAGATTGAGGAAAATAAGCCTTATT CTCTAAAGCGTGGGAAGCACGTGTATTCCATAAAGGCTAGaA ATGATGCCTTGGCAGAGTGTACAAGCAAATTTGTGACACAGT ATCCATGTATGATAAAAGGGTGTACTTCAGTCGTTACAAGTGA AAGCAATATCATCAGACATTATAAGTGTCATAAGTTGTCCAGG GCATTTACATCACAACACCGCAACATTCTTATTGTCTTTAAGC GATATGGCAACCCACAAGGAAAGGAAATCTCgGAGCAAGAAG ATGAAAAGAATGATAAGAAAGATCCTGATTCATCTGTcTTAGA GAAgAATGAcAACTCGGAACCAGCTGCcGCTCCACAGGAAGA AGGTAGAAAgGGTGAAAAGGATGAGATGGATGAGTTAACAGA ATTATTTATTACAAAGTTAATAAATGAAGACAGCACAAATGCA GAAAACCAAGGCAATACCACTTTAAAGGGAAATAACGAATTTC AGGAaCATGATTCCTGCACATCAGAAAGACAAAAGCCaGGTA ATTTaAAGAGAGTTTATAAgGAAAAgAACACTGTGCAGAGTAA GAAACGGAAGATTGATAAgACgGAGCCAGAAGTATCCTTGGT |

| | |
|---|---|
| | GGTAAATAATACACGGAAgGAGGAAGAGCCTGCCGTtGCAGT TCAGACCACTGAGGAGCATCCTGCATCCTTTGACTGGAGCTC CTTCAAGCCTATGGGATTTGAAGCATCaTTcCTGAAGTTTCTT GAAGAGTCTGCgGTGAAGCAGAAGAAAAATAGTGACAGAGAC CATTCAAACAGTGGAAGTAAAAGAGGATCCCATTCCAGCTCC AGAAGACATGTTGATAAGGCTGCTGTaGCTGGTAGCAGTCAT GTGTGTTCCTGTAAAGACAGTGAAATCTTTGTACAGTTTGCCA ACCCCTCAAAGCTTCAGTGCAGTGAGAATGTAAAAATTGTTTT AGACAAGACTCTTAAAGATCGCTCTGAGCTTGTCCTAAAACA GCTcCAGGAAATGAAACCTACTGTCAGTCTAAAAAAACTTGAA GTACTATCCAATAATCCAGATAGGACTGTTTTAAAAGAAATCA GTATAGGTAAAGCCACGGGCAGAGGGCAGTACTGA |
